# Multiple tasks viewed from the neural manifold: Stable control of varied behavior

**DOI:** 10.1101/176081

**Authors:** Juan A. Gallego, Matthew G. Perich, Stephanie N. Naufel, Christian Ethier, Sara A. Solla, Lee E. Miller

## Abstract

How do populations of cortical neurons have the flexibility to perform different functions? We investigated this question in primary motor cortex (M1), where populations of neurons are able to generate a rich repertoire of motor behaviors. We recorded neural activity while monkeys performed a variety of wrist and reach-to-grasp motor tasks, each requiring a different pattern of neural activity. We characterized the flexibility of M1 movement control by comparing the “neural modes” that capture covariation across neurons, believed to arise from network connectivity. We found large similarities in the structure of the neural modes across tasks, as well as striking similarities in their temporal activation dynamics. These similarities were only apparent at the population level. Moreover, a subset of these well-preserved modes captured a task-independent mapping onto muscle commands. We hypothesize that this system of flexibly combined, stable neural modes gives M1 the flexibility to generate our wide-ranging behavioral repertoire.

## INTRODUCTION

The generation of movement is crucial for survival. Whether seeking food, escaping a predator, or using tools to construct a shelter, motor behavior is arguably the ultimate purpose of the nervous system^1^. Primates, especially humans, have developed an advanced cerebral cortex that allows for a rich repertoire of arm and hand movements. The activity patterns of neurons in the primary motor cortex (M1) during such movements are accordingly complex; however, the mechanisms by which a single population of neurons can control varied behaviors remain unclear.

Historically, researchers have looked for reliable linear correlations between single neuron activity and specific movement parameters^2–5^. However, the wide variability of single neuron activity patterns^6,7^ has obscured the recognition of underlying principles. An intriguing alternative is that the computations mediating movement generation are performed at the population level, by interconnected cortical neurons^8–11^ whose coordinated activity commands the muscles that cause the behavior^10–13^. In this view, any correlates between single neuron activity and behavior are epiphemenonal^14,15^ and yield only a limited and distorted view of the causal relation between M1 and behavior.

We are currently able to monitor hundreds and even thousands of neurons simultaneously, a number that appears to be increasing exponentially^16^. Nonetheless, this is still a vanishingly small fraction of the number of neurons in motor cortex. To study neural function at the population level, we can describe neural activity in a high-dimensional *neural space* in which each axis represents the activity of one recorded neuron^9,17–19^. Even 10^2^ dimensions pose substantial theoretical and practical challenges; this space becomes unimaginably large as dimensionality increases to 10^5^ and beyond^20^. Fortunately, many dimensionality reduction studies^17,21^ within numerous brain areas show that neural activity explores only a limited, low-dimensional portion of the full neural space^9,22^. This sub-dimensional region, the *neural manifold*^9,18,19,23^, is determined by the explored neural population activity patterns (Fig. 1b). This neural manifold is spanned by *neural modes*, patterns of neural covariance that are thought to arise from the network connectivity^24^ (Fig. 1a) and thus unlikely to be voluntarily altered on a timescale of hours^18^.

**Figure 1.**
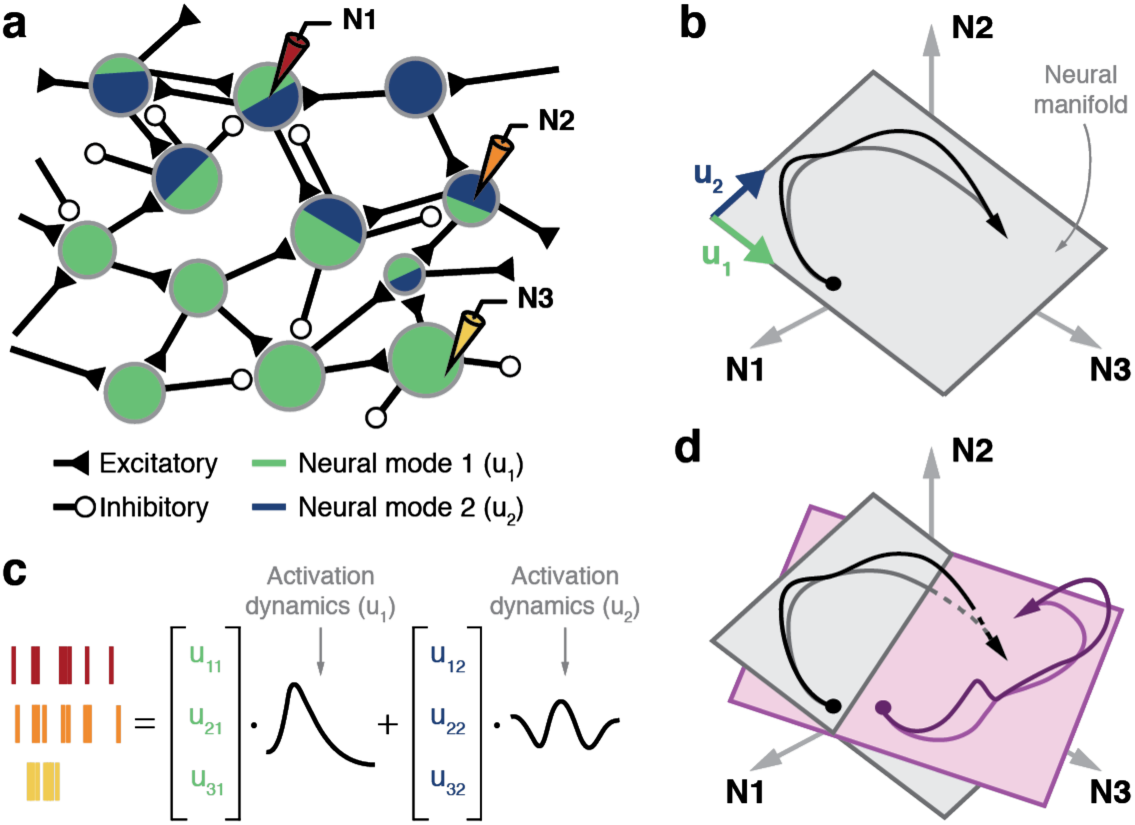
We hypothesize that different movement behaviors are caused by the flexible activation of combinations of neural modes. **(a)** The network connectivity within cortex results in the emergence of neural modes whose combined activation corresponds to specific activity patterns of the individual neurons. **(b)** Neural space for the activity patterns of the three neurons recorded in (a). The time-dependent population activity is represented by the trajectory in black (arrow indicates time direction). This trajectory is mostly confined to a two-dimensional neural manifold (gray plane) spanned by two neural modes (green and blue vectors). **(c)** The activity of each recorded neuron is a weighted combination of the time-varying activation of the neural modes. **(d)** Do neural manifolds for different tasks (show in gray and light purple) have similar orientations? Are the time-varying activations of the neural modes for two tasks (shown in black and purple) similar? These are the two critical questions that test our hypothesis.

We hypothesize that motor cortex generates varied behavior through the flexible activation of different combinations of neural modes. To examine this hypothesis, we identified M1 neural modes during a variety of distal limb tasks. Despite widespread differences in both neural activity and behavior, the structure of the neural modes was remarkably similar across different tasks. There were also striking similarities in the neural mode activation dynamics, the way these modes were recruited over time. Moreover, the *activation dynamics* of a specific subset of the neural modes were strongly predictive of muscle activation (EMG) patterns across the tasks; this indicates that these modes captured a task-independent component in the mapping from M1 to EMG. We thus propose that motor cortex generates different motor behaviors through the flexible activation of different combinations of fixed neural modes, and that the activity of single neurons simply reflects arbitrary one-dimensional samples of the manifold dynamics (Fig. 1c). The presence of neural modes in other brain areas (frontal^25^, prefrontal^26–29^, parietal^30,31^, visual^32–34^, auditory^35^, and olfactory^36^ cortices; see the reviews in Refs. 9,17) suggests that flexibly combined neural modes may be a general mechanism for neural computation.

## RESULTS

### Hypothesis, behavioral tasks and neural recordings

We addressed the hypothesis that motor behaviors are generated by the flexible activation of different combinations of neural modes by comparing both the structure of the modes identified during different motor tasks and their temporal activation dynamics. Our first prediction is that we will find similar modes across behaviors. Consider a simple three-neuron example (Fig. 1b). The population activity during movement traces a trajectory that could in principle explore any part of the neural space. In practice, correlations (covariation) between neurons constrain the population patterns and thus the region of neural space actually explored by the trajectory. If we use principal component analysis^17,23,37^ (PCA) to identify the dominant neural modes for this trajectory, we find two (***u***_*1*_ and ***u***_*2*_ in Fig. 1b). These two modes span a neural manifold^9,18,19^, the low-dimensional plane to which the trajectory is largely confined (Fig. 1b). We can assess the similarity of the neural modes for different tasks by comparing the orientation of the associated manifolds using principal angles^38^ (Fig 1d; details in later sections).

Our second prediction is that the time-varying activation of at least some neural modes (the *neural mode dynamics* or latent variables^9^) for different tasks will also be similar. This similarity follows from the assumed influence of network connectivity not only on the structure but also on the activation dynamics of the neural modes^39^. Canonical correlation analysis^40^ is a useful tool for performing this comparison^11^. It is crucial to note that this similarity could occur despite the high variability of single neuron activity across tasks.

To study M1 activity, we recorded data using 96-channel microelectrode arrays chronically implanted in the hand area of M1 of three rhesus macaque monkeys (monkeys C, T, and J). All surgical and behavioral procedures were approved by the Animal Care and Use Committee at Northwestern University. In each session, the monkeys performed one of two sets of motor tasks (Methods). The first set comprised several wrist tasks, including one-dimensional isometric, and both unloaded and elastic-loaded movement tasks^2,41,42^ (monkeys C and J); monkey J also performed a two-dimensional isometric task^43^. The other set of tasks (monkeys T and J) included a power grip task and a task that required a ball to be grasped, transported, and dropped^44^. One task of each set is illustrated in Fig. 2. In the one-dimensional isometric wrist task (Fig. 2a), the monkeys controlled cursor movements through the torque exerted at the wrist (Fig. 2b). In the power grip task^44^ (Fig. 2d), the monkeys reached for and grasped a pneumatic tube, squeezing it to achieve a target force (Fig. 2e). Successful completion of each task was associated with distinct patterns of muscle (Suppl. Fig. 1) and neural activity (Fig. 2c,f, Suppl. Fig. 2). The complex changes in neural activity patterns are quantified by the remarkably low cross-talk correlations (Fig. 2g).

**Figure 2.**
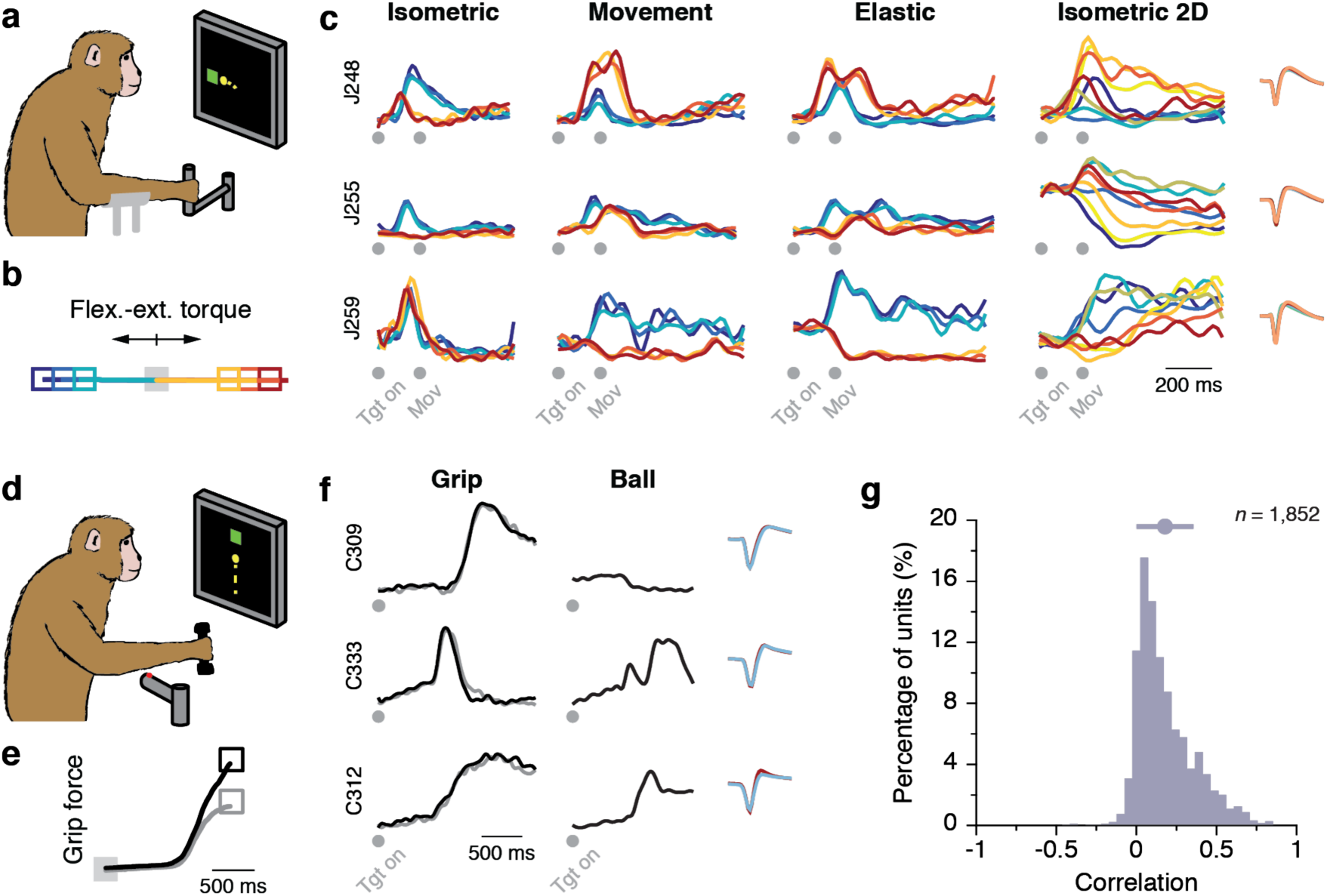
Tasks and recordings. **(a)** The wrist isometric center-out task. **(b)** Torque to acquire each target (colored squares). **(c)** Firing rates for three units illustrate the variety of observed activity patterns across units and how these patterns change across tasks. Right inset: action potential waveform for each task (each in a different color). Data for (b) and (c) are from monkey J, averaged over all the trials in one session, and colored according to target location (b; see also Suppl. Fig. 2). **(d)** The power grip task. **(e)** Grasp force trajectories to acquire each target (black or gray square). **(f)** Firing rates for three units show the variety of activity patterns observed during the grip task, and how they change in a complex manner for the ball task. Right inset: action potential waveform for each task (each in a different color). Data for (e) and (f) are from monkey C, averaged over all the trials in one session, and colored according to target (e). **(g)** Correlations between the activity pattern of each unit across each pair of tasks, pooled over all units, task comparisons, sessions, and monkeys; top error bar: mean ± s.d. correlation.

During each task we identified threshold crossings of both single- and multi-unit neural activity^18^ (number of units: 65.9 ± 16.9 across all datasets; mean ± s.d.; range, 45–91). For each task, we used PCA to identify the neural modes spanning a 12-dimensional (12D) neural manifold^9,17,23,37^ (Methods). Activity confined to these 12D manifolds accounted for at least 60% of the neural variance for all tasks, across all datasets (73.4 ± 6.5%; Suppl. Fig. 3b,c). This dimensionality is comparable to that reported for populations of neurons in the arm area of M1 during reaching^18,45^ and reach-to-grasp movements^46^. Notably, the majority of units contributed to all neural modes for all datasets, confirming that each neural mode captured a population-wide activity pattern (Suppl. Fig. 3a,d). The structure of the neural modes and their associated activation dynamics were robust against randomly sampled units; both were remarkably well preserved even if computed from only 50% of the recorded units (Suppl. Fig. 4). This observation further highlights that neural modes capture a physiological phenomenon that is shared among the entire population of neurons, and that the low-dimensionality of the neural manifold is intrinsic to the population activity and not an artifact of the small number of recorded neurons^22^.

**Figure 3.**
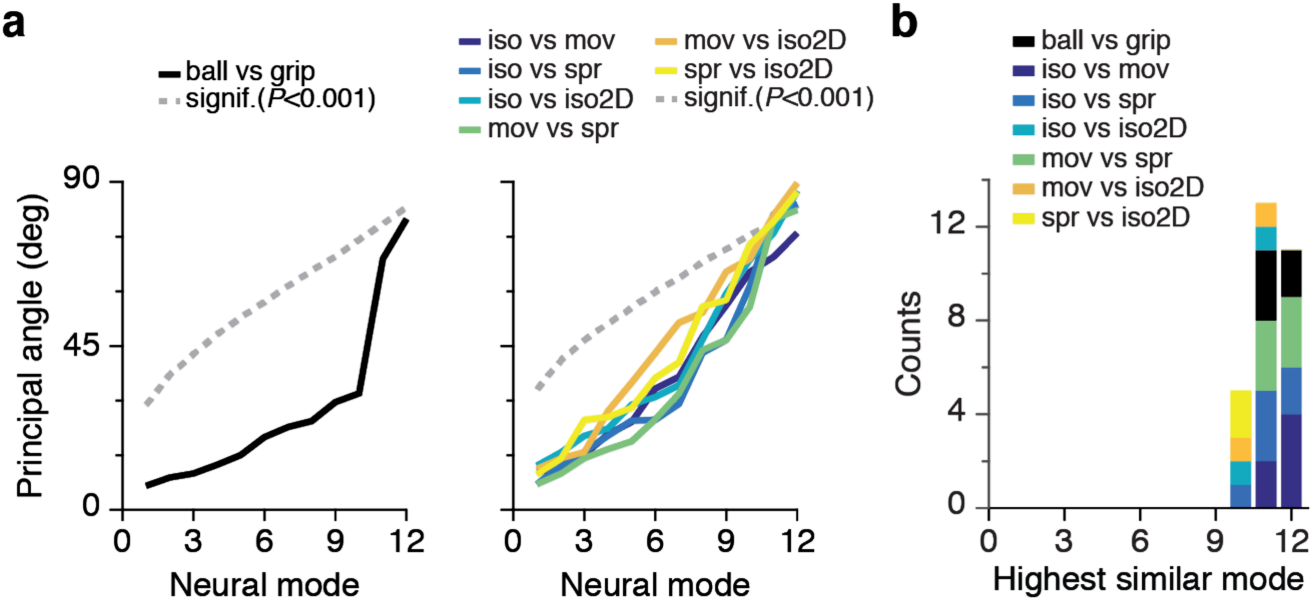
Principal angles between neural manifolds for two different tasks quantify the similarity of the corresponding neural modes. **(a)** Principal angles for one session of reaching and grasping tasks from monkey T (left) and for one session of wrist tasks from monkey J (right). Each pairwise comparison is shown as one colored trace (see legend). Leading principal angles were far below the *P*=0.001 significance level (dashed gray line), indicating similarities in the structure of the neural modes across tasks. **(b)** Number of neural modes for which all principal angles were significantly small across all monkeys and all pairs of tasks.

**Figure 4.**
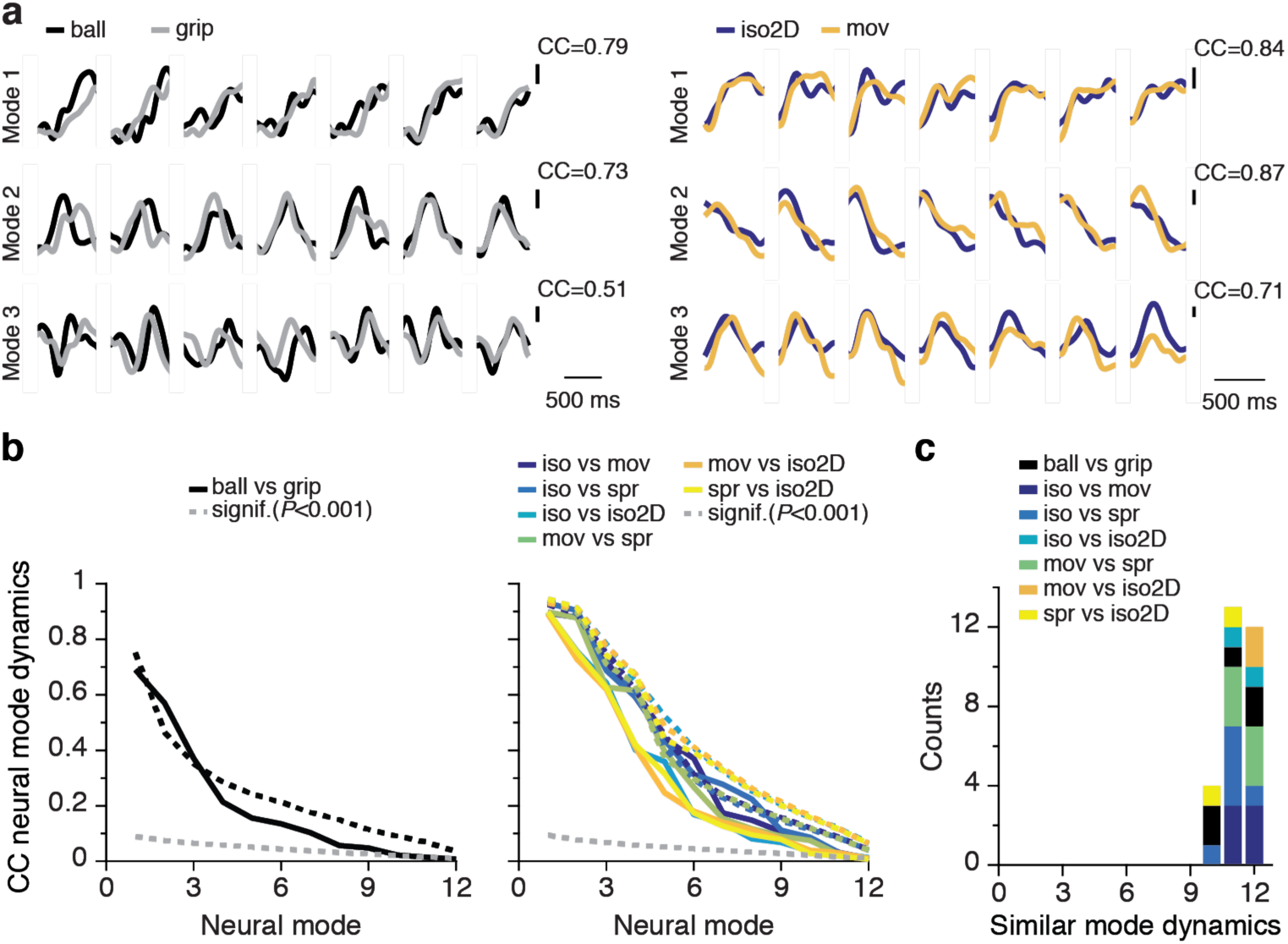
Canonical correlations between two different tasks quantify the similarity of the corresponding neural mode dynamics. **(a)** CCs for seven trials of the ball and power grip tasks for one session from monkey T (left), and for seven trials of one-dimensional movement and two-dimensional isometric wrist tasks for one session from monkey J (right). **(b)** CC between all pairs of reaching and grasping (left) and wrist tasks (right) in the same sessions as in (a) and (b). Each pairwise comparison is displayed as a color-coded solid line (see legend); the significance threshold (*P*<0.001) is shown as a dashed gray line. The dashed colored lines are upper bounds for across task comparisons provided by the average of the corresponding within task CCs (same color). **(c)** Number of significant CCs (*P*<0.001) across all monkeys and all pairs of tasks.

### Comparison of neural modes across motor tasks

To address our hypothesis that M1 generates movement through combinations of neural modes, we first tested whether the neural manifolds from different tasks were similarly oriented. To this end, we computed the 12 principal angles^38^ between the 12D manifolds for all pairs of tasks during each session (Methods). Our hypothesis predicts that these angles will be small. If M1 were able to recruit neurons in arbitrary combinations rather than as part of neural modes, the correlation structure would likely change across tasks, and the corresponding manifolds would differ significantly. Since developing an intuition for the expected value of the 12 principal angles between two 12D manifolds within a high-dimensional neural space is difficult, we interpreted our experimental results by comparing them to the distribution of angles obtained from a null hypothesis generated for pairs of randomly oriented planes (Methods, Suppl. Fig. 5). This distribution allowed us to set a very conservative threshold (*P*< 0.001) to consider the angles between these manifolds as significantly smaller than for randomly oriented ones (dashed grey lines in Fig. 3a).

**Figure 5.**
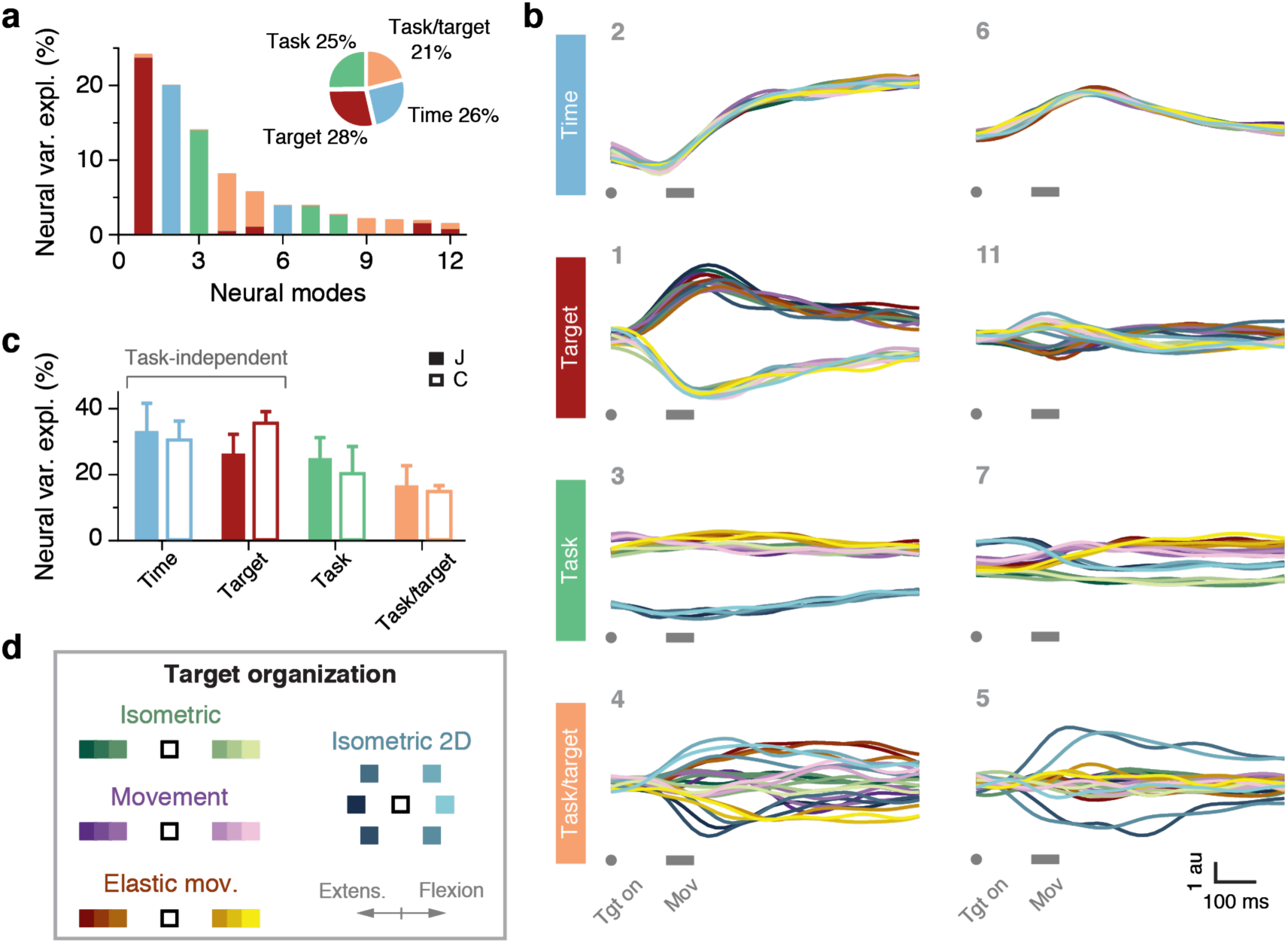
Task-specific and task-independent activation dynamics of the neural modes identified by dPCA. **(a)** Neural variance explained by each neural mode, and its relation to behavioral parameters for one example session from monkey J. Inset: amount of neural variance associated with each behavioral parameter across all twelve modes. **(b)** Activation dynamics of the eight leading neural modes, grouped in four sets based on the behavioral parameter they are most strongly associated with. The number on the top left of each panel indicates the ranking of that neural mode in terms of neural variance explained, as in (a). Each row corresponds to one behavioral parameter (vertical labels on the left). Each panel has 24 traces, corresponding to each of the 24 task/target combinations; color code shown in (d). **(c)** Total amount of neural variance explained for each behavioral parameter, averaged for all the sessions from monkeys J and C. Bars: mean + s.d. **(d)** Target locations for each task and color code for each task/target combination in (b). Each task is represented using a different color; extension targets are shown in dark colors and flexion targets in light colors

We found that the leading principal angles between task-specific manifolds were always very small, even with respect to our conservative threshold (examples in Fig. 3a). The three leading principal angles computed across all monkeys and pairs of tasks averaged 8.4 ± 2.3°, 11.3 ± 2.9°, and 15.1 ± 4.6°, all well below the chance level at *P*<0.001 (all datasets in Suppl. Fig. 6a). Even the tenth principal angle was always below this significance level (Fig. 3b). Therefore, as predicted by our hypothesis, there are strong similarities in the structure of the neural modes that span population activity during different motor tasks. The similarity implies that the structure of the population covariance patterns has a large fraction of preserved components (10 to 12 of the modes for the 12D manifolds), in contrast with the great variety of activity patterns that units exhibit across tasks (Fig. 2, Suppl. Fig. 2).

**Figure 6.**
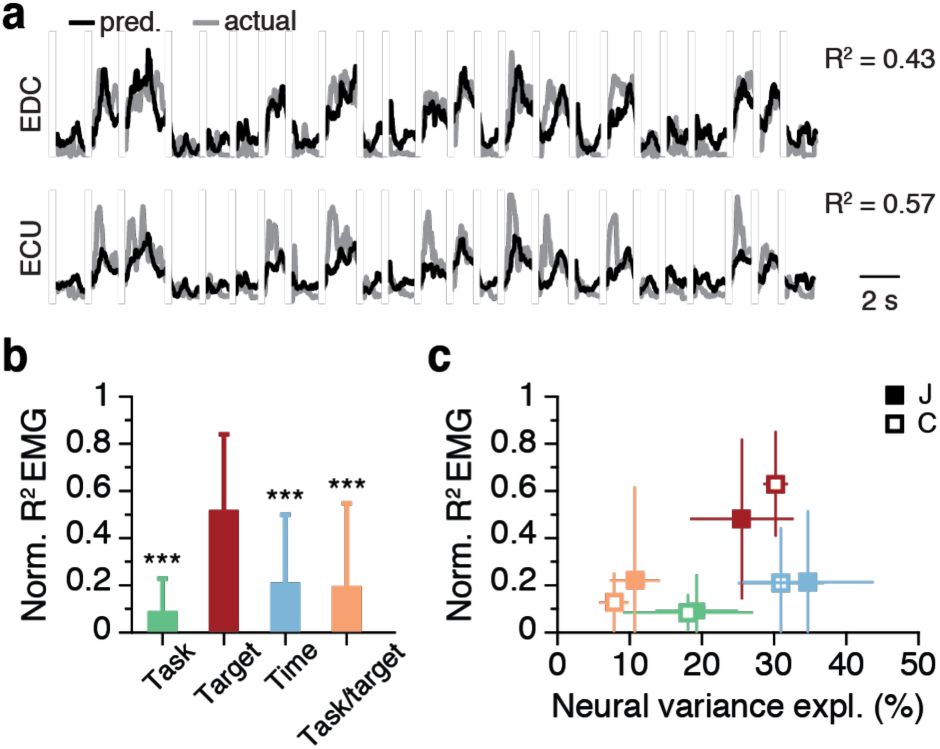
The activation of a subset of neural modes with task-independent dynamics captures a significant contribution to muscles commands. **(a)** Example EMG predictions for two muscles during 24 trials of the one-dimensional isometric task for one of the sessions from monkey J. Predictions were obtained with a decoder that took the activation dynamics of the two leading target-related neural modes as inputs. **(b)** Normalized R^2^ of the EMG predictions obtained from four types of decoders; each type took as inputs the activations of the two leading neural modes most strongly related to each of the four dPCA behavioral parameters. Performance was averaged over all muscles, tasks, and monkeys. Bars: mean + s.d; the *** denotes *P*∼ 0 (paired *t*-test). (c) Normalized R^2^ of the EMG predictions as a function of the neural variance explained by each of the four sets of neural modes identified with dPCA. Data for each monkey presented separately (legend). Squares: mean ± s.d; color code as in (b)

### Comparison of neural mode dynamics across motor tasks

Given the similarities in the structure of the neural modes of different tasks, we sought to understand whether their time-varying activation dynamics were also preserved. To address this question, we used canonical correlation analysis^11,40^ (CCA; see Methods). CCA is analogous to principal angles, but compares time-varying population signals, rather than the orientation of the neural manifolds that contain them. CCA yields as many canonical correlations (CC) as signals being compared; each CC ranges from 1 to 0, with 1 meaning perfect correlation.

Examples of CCs (Fig. 4a) reveal large similarities between the neural mode dynamics from different tasks. Across all monkeys and pairs of tasks, the three leading CCs were surprisingly high, averaging 0.85 ± 0.09, 0.72 ± 0.16, and 0.57 ± 0.17, respectively (examples in Fig. 4b; all datasets in Suppl. Fig. 6b). To develop a better intuition for these results, we compared these across-task CCs to the CCs for different trials of the same tasks (Methods). Strikingly, we found that the leading across-task CCs were quite similar to those of the within-task CCs, as indicated by how close the solid and dashed lines of the same color are for the leading dimensions (Fig. 4b: detailed results in Suppl. Fig. 7a,b). This result demonstrates the preservation of significant components of the neural mode dynamics across tasks. When pooling the results for all monkeys and all pairs of tasks, the leading ten CCs were always significantly above threshold (*P*<0.001 with bootstrapping; Fig. 4d). Interestingly, the leading CCs between neural mode dynamics were higher than the corresponding CCs between EMGs from different tasks (Suppl. Fig. 7c,d), suggesting that the well-preserved M1 activity cannot be trivially explained by similarities in muscle activation. Our finding that neural mode dynamics are better preserved across different tasks than is the activity of individual units supports the view that M1 may generate different behaviors by the flexible activation of different combinations of neural modes^8,9,18^.

### Task-specific and task-independent aspects of the neural mode dynamics

We have found that neural modes from different tasks are significantly aligned (Fig. 3) and that the leading neural mode dynamics are strongly correlated across tasks (Fig. 4). However, we also found differences in neural mode dynamics across the different tasks, as indicated by the monotonic decrease in CC: on average, the CC dropped below 0.3 when considering modes beyond the leading five. These results suggest an intriguing possibility: that the brain generates different behaviors by adjusting the activation of an existing set of neural modes in order to achieve the necessary motor output.

To investigate this possibility, we used demixed PCA (dPCA) to find a single neural manifold that spanned each set of tasks in a given session. We exploited dPCA’s ability to identify neural modes whose dynamics covary with specific behavioral parameters^36^ to understand the role of those modes with task-independent activation dynamics (Methods; Suppl. Fig. 8a). We looked for “time-related” modes whose activation dynamics depended only on time (i.e., progression through a trial), “target-related” modes that depended on the location of the target, “task-related” modes that depended only on the task being performed, and “task/target” modes that related to both task and target. Neural modes whose dynamics depend on time or target are task-independent, whereas neural modes whose dynamics depend on task or task/target interaction are task-dependent. We confined our analysis to the datasets for wrist tasks (one-dimensional isometric, movement, and elastic-loaded movement tasks for monkeys J and C, plus two-dimensional isometric for monkey J; see Methods) because of the similar spatial organization of the corresponding targets (Fig. 5d; Suppl. Fig. 1, 2).

As predicted by the small principal angles between tasks (Fig. 3, Suppl. Fig. 6a), dPCA found a single 12D neural manifold within which most of the population variance was contained (95.5 ± 0.5 % of the total variance accounted for by PCA, across all monkeys and sessions; Suppl. Fig. 8b). Each of the dPCA neural modes that spanned this manifold covaried almost exclusively with one of the chosen behavioral parameters (Fig. 5a, additional example in Suppl. Fig. 8c). More than half of the total neural variance (65.5% for monkey C, 59.0% for monkey J) was captured by neural modes that shared task-independent activation dynamics (Fig. 5c). This result illuminates the similarities in neural mode dynamics found with CCA.

Fig. 5b shows the dynamics of the eight leading neural modes for each of the 24 task-target combinations (four tasks × six targets; Fig. 5d) for one representative session (Suppl. Fig. 7d shows another session from a different monkey). The top row in Fig. 5b shows neural modes whose dynamics were virtually identical for all targets and tasks; their degree of similarity is striking given the different time courses of the corresponding motor outputs (Suppl. Fig. 1). The second row in Fig. 5b shows additional task-independent neural modes whose dynamics are related to the location of the target. These dynamics separated targets requiring wrist extension (dark colors) from those requiring wrist flexion (light colors), regardless of the specifics of the task.

The third row in Fig. 5b shows neural modes with task-specific dynamics that captured aspects unique to each task (Suppl. Fig. 1). For example, neural mode three is a task-related offset in the level of population activity that distinguishes the one-dimensional tasks from the two-dimensional task. This offset is present well before the movement is initiated, perhaps representing movement preparation^13,45,47^ that is task but not target specific. Finally, the neural modes shown in the fourth row in Fig. 5b covaried jointly with task and target, with complex activation patterns. Therefore, the observed similarities in neural mode dynamics, not apparent at the unit level (Fig. 2, Suppl. Fig. 2), are explained by the existence of neural modes with task-independent dynamics that account for a large percentage of the population variance.

### From neural modes to muscle commands

We have provided evidence that the neural population activity associated with different motor tasks can be generated through the activation of a small number of neural modes, some of which are task-independent. Given that M1 is the main cortical output to motoneurons^48,49^, it is reasonable to ask how the neural mode dynamics relate to the muscle commands that ultimately cause behavior. During any given task, muscle activity (EMG) can be reasonably well predicted as a readout from either the recorded population activity^12,50^ or the activation of the underlying neural modes^10,11,13^. Our dPCA analysis identified task-independent neural modes whose dynamics were explained by target location. The activation of these modes nicely separated wrist flexion from wrist extension, with largely task-independent temporal activation dynamics (Fig. 5b, modes 1 and 11; Suppl. Fig. 8c, modes 1 and 7). The existence of these neural modes suggests the possibility of a strong component of EMG activity that is also task-independent, in spite of the substantial variability observed in the time course of EMGs across the wrist tasks (Suppl. Fig. 1). Such EMG component should also follow from the target related neural modes in a task-independent manner. To test for the possibility, we built linear decoders^50^ that used the activation dynamics of the two leading target-related neural modes as predictors of the EMG activity generated during all the tasks in each session, including unloaded and spring-loaded movements as well as isometric contractions (Methods). If these neural modes captured a task-independent aspect of the M1 activity relevant to muscle activation, decoders taking them as inputs should yield good EMG predictions; also, these predictions should be better than those based on decoders that took the activation dynamics of the two leading neural modes from any other set (modes that depended on time, task, or task/target) as inputs.

For all monkeys, tasks, and muscles, decoders that took the two target-related neural modes as inputs made quite accurate EMG predictions (examples in Fig. 6a), with a cross-validated normalized R^2^ of 0.52 ± 0.32. These decoders thus were >50% as accurate as decoders that used all 12 neural modes as inputs (Methods); moreover, they far outperformed all other decoders based on modes related to any of the other three behavioral parameters (time, task, and task/target; Fig. 6b; by *t*-test, *P*∼0 for all three comparisons). Our ability to predict EMGs from the dynamics of the target-related neural modes could not be explained simply by the amount of variance that these modes captured (Fig. 6c). Specifically, the time-related neural modes accounted for more variance than the target-related neural modes, yet the corresponding EMG predictions were worse by a factor of ∼2.5. Thus, the target-related modes captured directions within the manifold that reflect a task-independent contribution of M1 activity onto muscle activity.

To further investigate the role of these target-related neural modes with fairly task-independent dynamics in the generation of EMGs, we examined the structure of the EMGs from all the tasks within a session using dPCA (Methods; Suppl. Fig. 9). This analysis revealed that half of the EMG variance was explained by *EMG modes*^51–53^ that were target-related (51.3 ± 5.0 %, across all datasets; Suppl. Fig. 9c). In spite of the different patterns of muscle activation required by unloaded movement and isometric contraction (Suppl. Fig. 1), the acquisition of similarly organized targets involved similar patterns of muscle co-activation (Suppl. Fig. 9a,b). We hypothesized that the prediction ability of target-related neural modes is due to their role in generating these target-related EMG modes. To verify this conjecture, we built another set of decoders that predicted only the target-related EMG modes from the four different sets of neural modes identified with dPCA. We found that these EMG modes were well predicted as the readout of the target-related neural modes (normalized R^2^: 75.6 ± 19.1 %; Suppl. Fig. 9d). We have thus identified a stable, task-independent component in the mapping from specific neural modes to specific EMG modes.

## DISCUSSION

Many prior studies have tried to understand how M1 controls movement by looking for a fixed relationship between single neuron activity and behavioral parameters (what neurons *encode* or *represent*). Critically, several of these studies appear to be contradictory, others suggest that different subclasses of neurons encode different behavioral parameters, and others that what a given neuron encodes changes across behaviors^4,7,14^. In light of these results, several groups have begun to investigate how the coordinated activity of populations of neurons relates to motor performance^14^. In particular, there has been interest in the dynamics of population activity within the neural manifold^8,9,45^. Thus far nearly all these manifold studies investigated manifolds associated with the execution of a single task, except for a recent study comparing reaching and walking in mice^54^. Here, we present the first comparison of neural manifolds computed across as many as six different upper limb skilled tasks. Do neural manifolds for different tasks have similar orientations? Are the time-varying activations of the neural modes for different tasks similar? These are the two critical questions that we investigate.

Our results show that motor behaviors may be generated by flexibly combining the activation of fixed neural modes^9^. For a variety of upper limb motor tasks that required quite distinct patterns of neural activity, the structure of the neural modes was largely preserved. Moreover, the activation dynamics of some neural modes were also strikingly correlated across different tasks. A subset of these neural modes that had predominantly task-independent dynamics captured a consistent mapping onto task-independent components of muscle activity. These results provide new insight into how movement is generated by the motor cortex, and suggest that cortical circuits with fixed connectivity may perform different functions through the flexible activation of different combinations of acquired neural modes.

There is increasing evidence that neural covariation patterns captured by the neural modes arise from the underlying network connectivity. In cat primary visual cortex (V1), optical imaging has revealed that the correlation between a neuron’s spontaneous activity and the state of its surrounding population is very similar to the corresponding correlation during stimulus-evoked activity^55^. This remarkable finding was later replicated in similar recordings from mouse V1^24^, in a study that also showed that each neuron’s weighted contribution to the leading neural mode correlated with the number of synapses onto that neuron. In addition, the response of a given neuron to optogenetic stimulation of the population could be predicted by the correlation between the spontaneous firing rate of that neuron and that of the population. These results provide strong albeit indirect evidence that network connectivity may largely determine the observed neural correlations, and thus the structure of the neural modes. For M1, the most convincing evidence relating neural modes to network connectivity comes from a brain-computer interface study in which monkeys attempted to produce altered neural covariance patterns^18^. The generation of new covariation patterns (new neural modes) during a single session was considerably more challenging than activating the existing neural modes in novel combinations.

To help understand how the activity of populations of M1 neurons varies so as to control movement during different upper limb behaviors, we searched for commonalities in the neural modes across different tasks^56^. Prior studies have shown that single neuron activity changes in complex ways across tasks, and that the variables that those neurons represent also change^4, 7, 14, 57–59^ (Fig. 2, Suppl. Fig 2). In contrast, the results we report here demonstrate that the structure of the neural modes is well-preserved (Fig. 3), and so are the activation dynamics of some of them (Fig. 4). The well-preserved structure of the neural modes is consistent with previous findings that relate network connectivity to the neural modes that span the manifold^18, 24^. If the neurons in a population are involved in causing upper limb movement during different learned behaviors, and if the connectivity among these neurons determines the structure of the neural modes, then it is reasonable to expect similarities across manifolds for the different tasks.

Intriguingly, a subset of the well-preserved neural modes had activation dynamics that were virtually the same regardless of the task or the movement generated to reach a specific target –the “time-related” modes, top row in Fig. 5b. What is the potential role of these modes? Neural population activity – and thus the activation of neural modes – reflects the population’s response to inputs, its internal computations, and its outputs^11^. Neural modes with task- and target-independent activation dynamics are unlikely to reflect population inputs or outputs, as these should differ across movements and behaviors. Instead, they probably capture internal computations. The role of such computations is unclear, but in this case the dynamics of these modes may reflect the population switching to an M1 *movement state*, as suggested by the observation that the activation dynamics of the leading time-related mode during an instructed delay reaching task predicted reaction time with great accuracy^60^. Other computations that these time-related modes could capture are switching from a “postural control” or “holding still” mode to a “movement control” mode^61, 62^, or modulating spinal reflexes prior to movement onset^63^.

Given the extensive connections of M1 to motoneurons and spinal interneurons^48,49^, we expect that readouts of the neural mode dynamics will map onto muscle commands^10,13^ (EMG). Here, we found a specific subset of neural modes with a task-independent mapping to EMGs (Fig. 6). Such task-independent mapping might simplify limb control. For example, rapid motor adaptation to a force field perturbation appears to be accomplished by exploring alternative neural mode dynamics only within the manifold that controls the unperturbed movement^64^. Similarly, corrective movements in response to visual perturbations are driven by mode dynamics confined to specific dimensions of the unperturbed manifold^65^. Either strategy would likely become much harder to carry out using a condition-dependent mapping onto muscle commands, because the brain would need to rapidly modify both neural activity within M1 and how it gets projected onto muscle activity.

The existence of a task-independent component of the M1 to EMG map might be due to a degree of similarity in the target structure across wrist tasks considered here; this similar organization probably causes the observed task-independent component in muscle co-activation patterns (Suppl. Fig. 9). Indeed, our results are in contrast to the behavior-specific mapping found in mice when forelimb M1 population activity during reaching was compared to that during treadmill walking^54^. In the comparative analysis of these two tasks, the corresponding manifolds were found to be orthogonal^54^ and exhibited none of the structural and dynamical similarities found here. This lack of similarity is likely due to M1 being less directly involved in the control of treadmill walking than of reaching^54,66^. Even if the similarities reported here are to some extent induced by similarities among tasks, it is still remarkable that an M1 to EMG mapping based on the distinction between flexor and extensor muscle activation will be preserved across isometric and movement tasks.

Neural modes are likely not restricted to motor cortices: evidence of them has been found in visual^32–34^, olfactory^36^, auditory^35^, frontal^25^, prefrontal^26–29^ and parietal^30,31^ cortices (see Refs 9,17 for recent reviews). These observations raise an intriguing question: do populations of neurons in these areas also use a flexible activation of neural modes to perform different functions? The application of dPCA to neural recordings during sensory discrimination tasks revealed neural manifolds with stimulus-related, decision-related, and time-related modes, in both monkey prefrontal and rat olfactory cortices^36^. Moreover, during a working memory task, population activity in prefrontal cortex was associated with a manifold spanned by modes that related linearly to memory storage and stimulus response^27, 66^. Thus, population activity in multiple cortical areas is associated with neural manifolds whose modes relate strongly (and linearly) to task-relevant parameters. The similarity between these results and the ones reported here for M1 (Fig. 5) suggests that populations of neurons in other brain areas could perform a variety of their specific functions by activating different combinations of neural modes, in a manner similar to how M1 seems to control different upper limb movements.

A potential limitation of our study is the attempt to infer general motor control principles based on the analysis of stereotypical laboratory tasks. We found consistent results from two groups of six different tasks (wrist and reach-to-grasp datasets; Figs. 2-4), which suggests that our results are not a simple consequence of comparing two overly similar behaviors. An interesting extension of the present work would be to study the structure and activation dynamics of the neural modes during more complex natural behaviors involving upper limb use. The first question would then be that of the dimensionality of the resulting M1 manifold. For standard laboratory motor tasks, M1 manifold dimensionality appears to be under ten (Suppl. Fig. 3; Refs. 10, 13, 18,45–47,64,65, and the studies discussed in Refs. 9,22). However, theoretical derivations show that manifold dimensionality increases with task complexity^22^. An increase in manifold dimensionality would require an increase in the number of recorded neurons in order to reliably map the manifold^22^. Therefore, it may be the case that we do not yet have the technical means to record from enough neurons to map the M1 manifold associated with unconstrained behaviors.

Notably, the dimensionality of the neural manifolds in motor cortical areas may decrease when getting closer to the main output in M1. For a standard center-out reaching task, the dimensionality of the M1 manifold was considerably smaller (almost half) than that of the manifold in the upstream dorsal premotor cortex^64^ (PMd). Since PMd is involved in integrating inputs from several areas and forming a motor plan that is then projected to M1^67^, it seems reasonable that its population neural activity will be more complex than that of M1 and thus be associated with a higher dimensional manifold. Another aspect that may impact the dimensionality of the M1 manifold is that an upper bound may be imposed by the intrinsic dimensionality of the limb dynamics.

In summary, we have shown that the manifolds associated with neural population activity during different motor tasks have similar orientation. Moreover, the activation dynamics of some of the spanning neural modes are also strikingly correlated across behaviors, in contrast with the highly varied patterns of muscle and neural activity. These results support the notion that motor cortex may control movement during different behaviors through the flexible activation of different combinations of neural modes, neural covariation patterns that reflect network connectivity. We further suggest that a similar mechanism may underlie the ability of other cortical areas to perform a wide variety of non-motor functions.

## Acknowledgements

This work was supported in part by Grant FP7-PEOPLE-2013-IOF-627384 from the Commission of the European Union (J.A.G.), by Grant F31-NS092356 from the National Institute of Neurological Disorder and Stroke and Grant T32-HD07418 from the National Center for Medical Rehabilitation Research (M.G.P.), by Grant DGE-1324585 from the National Science Foundation (S.N.N.), by Grant 22343 from the Fonds de Recherche du Québec–Santé (C.E.), by Grant NS094748 from the National Institute of Neurological Disorder and Stroke (S.A.S.), and by Grant NS053603 from the National Institute of Neurological Disorder and Stroke (L.E.M.).

## METHODS

### Experimental subjects

We recorded data from three 9-10 kg male *macaca mulatta* monkeys (J, C, T) while they performed one of two sets of wrist or reach-to-grasp motor tasks over several sessions (see Tasks, below). The monkeys were implanted with a 96-channel microelectrode silicon array (Utah electrode arrays, Blackrock Microsystems, Salt Lake City, UT) in the hand area of M1, which we identified intraoperatively through microstimulation of the cortical surface. For monkey C, we recorded neural activity for each of the two sets of tasks using different microelectrode arrays, which were sequentially implanted in a different brain hemisphere. The monkeys were also implanted with intramuscular EMG electrodes in a variety of wrist and hand muscles. We report data from the following muscles: Monkey J: flexor carpi radialis (FCR), flexor carpi ulnaris (FCU), extensor carpi radialis (ECR), extensor carpi ulnaris (ECU), flexor digitorum profundus (FDP), flexor digitorum superficialis (FDS), extensor digitorum communis (EDC; radial and ulnar aspects), brachioradialis, and supinator; Monkey C: FCR, FCU, ECR, ECU, FDP (radial and ulnar aspects), FDS (radial and ulnar aspects), EDC (radial and ulnar aspects), flexor pollicis brevis (FPB), opponens pollicis, and extensor pollicis longus; Monkey T: ECR, ECU, FCR, FCU, FDP (radial and ulnar aspects), FDS (radial and ulnar aspects), EDC, FPB, first dorsal interosseous (FDI). For the wrist tasks of monkey C, we recorded EMGs using pairs of gelled surface electrodes placed over FCR, FCU, ECR, ECU, FDS and EDC. Additional details about the surgical methods and postoperative care can be found in our previous publications^50, 43^.

### Tasks and recordings

In each session, monkeys performed either a set of reach-to-grasp tasks, or a set of wrist tasks (Fig. 2). All monkeys had been trained prior to their implant surgeries, and were proficient at the tasks at the time of the recordings. Monkeys C and T performed the set of reach-to-grasp tasks, which comprised the “ball” and power “grip” tasks (monkey C, three sessions; monkey T, two sessions). In the ball task, monkeys had to reach to a ball (diameter 24, 35, or 40 mm), grasp it, and then transport it and drop it in an open cylindrical container^44^. In the power grip task, monkeys reached to and grasped a pneumatic tube that then had to be squeezed to control the movement of a cursor used to acquire one of two or three one-dimensional force targets^44^. Monkeys initiated both tasks by resting their hand on a touch pad, and waited for a target (or go signal, for the ball task) to be presented. Monkeys C and J performed the wrist tasks, which comprised three one-dimensional tasks^2, 41, 42^: an isometric task, a movement task, and an elastic loaded movement task (both monkeys, three sessions); monkey J also performed a two-dimensional isometric center-out task^43^ in two of three sessions (see Fig. 2a, b). Throughout the paper, we abbreviate these tasks “iso,” “mov,” “spr,” and “iso2D,” respectively. As for the reach-to-grasp tasks, monkeys could initiate movement after the target was presented. During the experiments, we recorded neural and EMG data, as well as kinematics or force, depending on the task. All data were saved to disk and analyzed in Matlab (The Mathworks Inc., Natick MA) using purposely-written scripts; for the demixed principal component analysis (see below), we used the publically available toolbox from the Machens lab^36^ (https://github.com/machenslab/dPCA).

To characterize neural population activity, we identified threshold crossings from each electrode, which included well-discriminated single-unit as well as multi-unit activity. Throughout this paper we refer to these as *units*, without distinction. For each session, data included all units whose average waveform, triggered by the threshold crossing, remained stable across all tasks (examples in Fig. 2, Suppl. Fig. 2). We did not choose neurons based on tuning, modulation depth, or any other property. To obtain a smooth discharge rate as function of time, we applied a Gaussian kernel (s.d.: 50 ms) to the binned square-root-transformed firings (bin size: 20 ms) of each unit^23^.

The EMG envelope, a proxy for the neural commands to muscles, was computed by a sequence of high-pass filtering (4^th^ order zero-phase Butterworth filter, *f*_*c*_: 10 Hz), rectification, and low pass filtering (4^th^ order zero-phase Butterworth filter, *f*_*c*_: 50 Hz) of the raw EMG signals. We subsequently normalized these EMG envelopes by the 99^th^ percentile of their distribution across all tasks for each given session. We used single-trial data for all the analyses except for dPCA, a method that requires trial-averaged data^36^ (see details below). A trial was defined from target presentation until the monkey received a reward; the very few unsuccessful trials were discarded. For trial averaging, we computed the mean firing rate (peristimulus time histogram) from target presentation until an end time determined by the shortest time to reward. We used both the reach-to-grasp and wrist datasets for all the analyses except for the dPCA; the latter requires target equalization across tasks, which can only be achieved for the wrist tasks (see main text and dPCA section below). In every session, we compared tasks across all possible pairs.

### Task-specific neural manifolds and neural mode dynamics

The activity of *n* recorded units was represented in a *neural space*, an *n*-dimensional sampling of the state of M1. In this space, the position along each axis represents the firing rate of the corresponding unit (Fig. 1b). Within this space, we computed the low-dimensional *neural manifold* associated with each task by applying principal component analysis (PCA) to the smoothed firing rates of all *n* units for that task. PCA finds *n* principal components (PCs), each a linear combination of the firing rates of the units that maximizes the amount of shared variance (covariance). The PCs are ranked according to the amount of variance in the original data that each explains. We defined *m*-dimensional task-specific manifolds that accounted for most of the neural population variance by keeping only the leading *m* PCs (Fig. 1b). We chose *m*=12, to account for at least 60% of the total neural variance for all tasks and monkeys (Suppl. Fig. 3). Importantly, the results were not sensitive to the manifold dimensionality, as previously reported^18,45,47^. Each PC is a neural mode, a specific direction within the manifold; together, the neural modes provide a basis that spans the neural manifold. We computed the *neural mode dynamics* by projecting the *n*-dimensional, time-varying neural population activity onto each of the *m* neural modes (PCs) of the neural manifold.

### Comparison of task-specific neural manifolds

Principal angles provide a measure of the relative alignment of two *m*-dimensional manifolds in terms of the *m* angles between sequentially aligned pairs of basis vectors^38^. These vectors, selected in each manifold so as to systematically minimize the angle between them, provide a new basis in each of the two manifolds being compared. Note that manifold directions chosen to minimize the angles between manifolds are not necessarily those that maximize variance within each of the two manifolds; it is thus not the angles between the PC neural modes that determine the principal angles. Our hypothesis that task-specific manifolds are similar implies that the leading principal angles will be small.

To compute the principal angles between two *m*-dimensional manifolds *A* and *B* embedded in an *n*-dimensional neural space, we follow the method by Björck and Golub^38^: consider the corresponding bases *W*_*A*_ and *W*_*B*_ provided by the PC neural modes, construct their inner product matrix, and perform a singular value decomposition to obtain

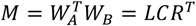

Here *W*_*i*_, *i* = *A*, *B* are the n by m matrices that define the task-specific manifolds *A* and *B*; the corresponding PC neural modes are their column vectors. The matrix *c* is a diagonal matrix whose elements are the ranked cosines of the principal angles *θ*_2_, *i* = 1… *m*:

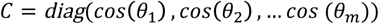

Note that by construction, the principal angles are ordered form smallest to largest.

To assess whether the experimentally obtained principal angles between pairs of task-specific manifolds were small, we compared them to empirically generated distributions of principal angles (example distributions in Suppl. Fig. 5a). We obtained these distributions by computing the principal angles between 10,000 pairs of randomly generated 12D manifolds embedded in spaces with dimensionality equal to that of each of the datasets we studied. We used the 0.1^th^ percentile of those distributions to define a stringent threshold below which angles can be considered significantly small (with a probability *P*< 0.001). As shown in Suppl. Fig. 5b, the threshold angles between 12D manifolds increased with the dimensionality of the neural space.

### Comparison of task-specific neural mode dynamics

To investigate potential similarities in neural mode dynamics across tasks, we compared the corresponding task-specific manifold dynamics using *canonical correlation analysis* (CCA). The method systematically finds new directions within each manifold such that the corresponding one-dimensional projected dynamics are maximally correlated. As is the case with the manifold directions used to compute principal angles, these directions are not necessarily those of the PC neural modes selected to maximize projected variance.

Consider again the two manifolds *A*, *B* to be compared. We start by projecting the dynamics of each manifold in this pair onto the corresponding PC neural modes, to obtain two *T* by m latent variables matrices *L*_*A*_ and *L*_*B*_; here *T* is time duration of all concatenated trials for a given task. The CCA finds two linear transformation matrices, one for each of the two mode dynamics matrices *L*_*i*_*, i* = *A*, *B*, to obtain new directions within the manifolds so that the dynamics projected onto these new directions within each manifold are maximally correlated^40^.

The CCA starts with a QR decomposition of the latent variables matrices *L*_*A*_ and *L*_*B*_, *L*_*A*_ = *Q*_*A*_*R*_*A*_, *L*_*B*_ = *Q*_*B*_*R*_*B*_ The first m column vectors of *Q*_*i*_,*i* = *A*, *B* provide a basis for the column vectors of *L*_*i*_,*i* = *A*, *B*. We then construct the inner product matrix of *Q*_*A*_ and *Q*_*B*_ and perform a singular valuen decomposition of the inner product matrix to obtain

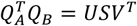

The elements of the diagonal matrix *S* are the canonical correlations (CCs). As for principal angles, the canonical correlations are by construction sorted from largest to smallest.

For this analysis, the matrices *L*_*i*_,*i* = *A*, *B* included all the concatenated trials for each of hose two tasks. To assemble these data matrices, we first equalized the number of trials across all the tasks within the corresponding session. For each trial, we used either a 700 ms long (wrist tasks) or a 1,000 ms long (reach-to-grasp tasks) window of neural data, starting around target onset. When comparing two one-dimensional wrists tasks, we matched the trials by target location; when comparing the two-dimensional isometric task to any of the one-dimensional tasks, we distributed the trials to vertical targets in the former task evenly across trials to each of the targets in the one-dimensional task. No target-matching was done for the reach-to-grasp tasks, as the ball task had no targets. We did not exclude trials based on their execution time, or based on the EMG, kinematics, or force patterns.

We used an analysis of inter-trial variability for each task to obtain an upper bound for the across-task CCs. To compute this upper bound, we first computed within-task CCs by dividing all the trials for a given task into two random target-matched subgroups (100 repetitions), and calculated the corresponding CC. We used the 99.9^th^ percentile of each within-task CC distribution as the upper bound CC value, and obtained an across-task upper bound for each pair of tasks by averaging the upper bounds of the two corresponding tasks. Actual across-task CC values close to this upper bound indicate remarkably similar neural mode dynamics, with differences comparable to those expected from within-task fluctuations. We also used bootstrapping to assess the significance of the across-task comparisons of neural mode dynamics (10,000 shuffles over time of one of the two sets of mode dynamics being compared); the 99.9^th^ percentile was again used as significance threshold (*P*< 0.001).

### Identification of task-independent and task-specific neural mode dynamics

To understand the role of these preserved neural mode dynamics in movement generation, we used another linear dimensionality reduction method, *demixed PCA*^36^ (dPCA). This approach identifies a single neural manifold for all the data (here, for all tasks), spanned by neural modes whose dynamics are linear readouts of the dynamics associated with chosen behavioral parameters^36^. The ability to find a single neural manifold for all the tasks that we examined is due to the strong similarity in the orientation of the corresponding task-specific manifolds (Fig.3)

Mathematically, dPCA represents the mean-subtracted, trial-averaged activity from all units concatenated over all tasks and targets within a session as a neural data matrix *X*. This matrix is decomposed as a sum of activities *X*_ø_ each related to a specific behavioral parameter ø. Thus, *X* is written in terms of the usually called marginalizations *X*_ø_ and the trial-to-trial-noise *X* _*noise*_:

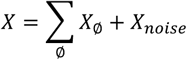

The marginalization ensures that the *X*_ø_ are uncorrelated, and that the *n* by *n* covariance matrix *c* = *XX*^*T*^. is the sum of covariance matrices, one for each marginalization:

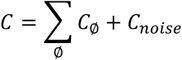

Dimensionality reduction in dPCA is based on the minimization of a reconstruction error

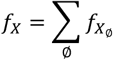

with

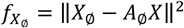

The minimization of the reconstruction error becomes equivalent to a classical regression problem with ordinary least squares solution^36^:

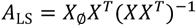

In dPCA, the experimenter chooses the rank *m* of the *n* by *n* matrix *A*; m is the dimensionality of the manifold. The ordinary least square problem thus becomes a reduced-rank regression problem that is solved using singular value decomposition. A detailed description of dPCA for neural population data has given by Kobak, Brendel and colleagues^36^; notably, this implementation of dPCA has an analytic as opposed to numerical solution.

The behavioral parameters ø used here are: *time* along the trial, *task, target location*, and the combination *task/target location*. We performed the dPCA analysis on the wrist tasks, as these datasets included three or four tasks for which six targets were similarly located in space (see target organization in Suppl. Fig. 1, 2). We equalized the number of trials across targets and tasks. As for the previous analyses, the chosen manifold dimensionality was *m* = 12. In spite of the constraint that the time-varying activation of each neural mode has to covary with one or a few of the chosen behavioral parameters, the neural variance explained by the neural modes identified with dPCA was very similar to the variance explained by the PCA modes (see example in Suppl. Fig. 8b).

### Relationship between the neural mode dynamics and EMGs

To understand the role of the neural modes in movement generation, we investigated how their dynamics related to the ongoing muscle commands (EMGs) by building standard linear decoders as previously used by our group^50, 68^. We were particularly interested in the role of the target-related but task-independent neural modes identified with dPCA. To assess whether these target-related modes captured a constant (task-independent) component in the time-dependent muscle activations (EMGs), we compared the predictions of decoders that used the dynamics of target-related modes as inputs to the predictions of decoders that used as inputs the dynamics of the other three sets of modes (time-related, task-related, and task-target-related).

The neural to EMG decoders were multiple-input single-output linear filters followed by a static non-linearity:

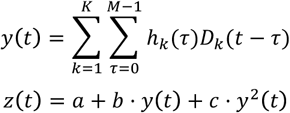

Where *z*(*t*) is the predicted EMG, obtained by applying a static non-linearity to the output *y t* of the linear model. The linear model estimated the EMG as a linear combination of the current and past values of the neural mode dynamics, *D*_*k*_, *k* = 1,2, weighed by the coefficients *h*_*k*_ (τ), where τ represents time into the past. The filter coefficients *h*_*k*_ (τ) were obtained using the autocorrelation and crosscorrelation matrices of the decoder inputs and outputs^50^. The coefficients *a*, *b* and *c* of the second order polynomial in the static non-linearity were computed using least squares error minimization.

We built a single decoder for each behavioral parameter ø using data from all the tasks that the monkeys performed during one session. We assessed the quality of fit on single trial data in terms of the normalized coefficient of determination (*R*^2^), which is the ratio of the *R*^2^ of the EMG Predictions based only on the activation dynamics of the neural modes related to a specific behavioral parameter ∅, to the *R*^2^ of the EMG predictions based on the activation dynamics of all 12 neural modes. Fits were cross-validated (30 s folds) in all cases. We compared EMG predictions across marginalizations using a paired *t*-test including each fold.

To interpret our decoding results, we decomposed the EMGs from all the tasks within a session into *EMG modes* using dPCA. We followed the same methods as for the neural data, and chose *m* = 4 modes, as this value maximized the EMG variance explained with dPCA across all datasets. When predicting subsets of EMG modes, we used decoders with the same structure described above, and followed the same cross-validation procedure.

### Control analyses

To probe the dependence of manifold geometry and neural mode dynamics on the dimensionality of the embedding neural space, we performed unit-dropping numerical experiments. We first tested whether the observed geometry of the manifold depended on details of the activity of recorded units. To this end, we selected two random subsets of all recorded units, obtained the 12D manifolds spanned by the 12 leading PC neural modes, and computed the principal angles between them. We repeated this operation dropping 10, 20, 30, 40, and 50% of all recorded units (100 random pairs in each case). If manifold geometry was invariant under choice of units, these principal angles should be small (see Suppl. Fig. 4a). We also tested whether the observed neural mode dynamics depended on details of the activity of recorded units. For this analysis, we selected a random subset of all recorded units, obtained the 12D manifold spanned by the 12 leading PC neural modes, and then projected the population activity onto these neural modes to obtain their activation dynamics. We then used CCA to compare the dynamics of these modes to the dynamics of the 12 leading modes computed from all recorded units. We repeated this operation dropping 10, 20, 30, 40, and 50% of all recorded units (100 random pairs in each case). If neural mode dynamics did not depend on the specific choice of units, the leading CCs should be close to 1 (see Suppl. Fig. 4b).

To assess the similarity of neural mode dynamics across tasks, we obtained a within-task upper bound to the maximum expected across-task similarity in neural mode dynamics (see details above). To quantify how close the across-task CCs came to the correspondingly averaged within-task upper bounds, we computed their ratio to obtain a 12-point function for each task comparison (see Suppl. Fig. 7a,b).

To monitor changes in the relation between neural mode dynamics and EMGs for different tasks, we first assessed the across-task stability of the EMGs. We applied CCA to the muscle activation patterns for each pair of tasks, using the same methods as for the across-task comparison of neural mode dynamics. To quantify the across-task stability of mode dynamics, we computed the ratio of the across-task CC in neural mode dynamics to the across-task CC in EMGs (see Suppl. Fig. 7c,d), for as many dimensions as EMGs we had available (we typically had less than 12 well-recorded muscles in any given session).

**Supplementary Figure1.**
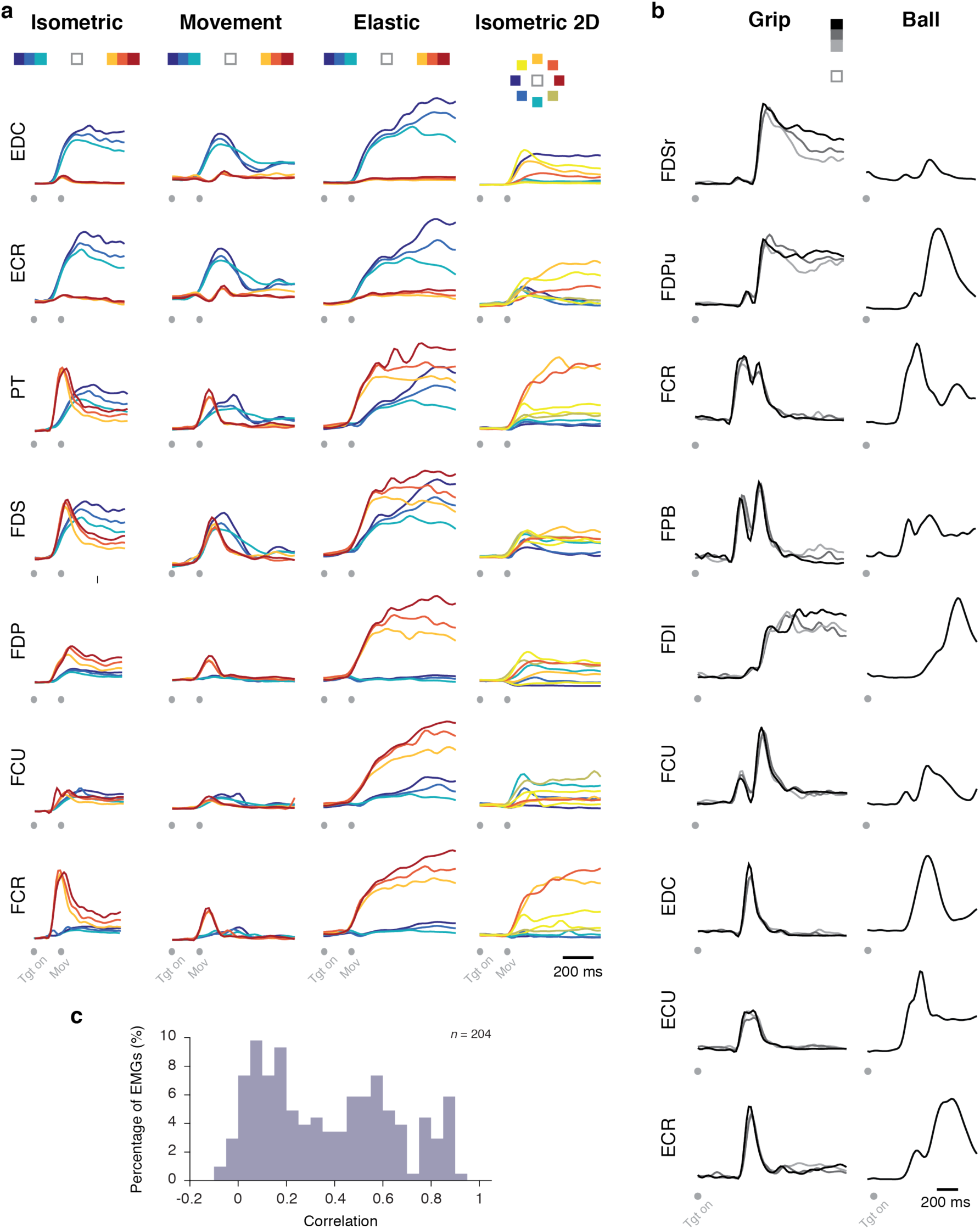
Examples of muscle activity patterns (EMGs) illustrate their broad diversity both across tasks and across targets for a given task. **(a)** EMG envelope of seven wrist and hand muscles for one session in which monkey J performed all four wrist tasks. EMGs are colored according to target location (below task name). **(b)** EMG envelope of nine wrist and hand muscles for one session in which monkey T performed the two reach-to-grasp tasks. Data organized as in (a). **(c)** Correlation of the activity of each muscle across two different tasks, pooled over all muscles, tasks, sessions, and monkeys. Note the strong presences of low correlation instances. Muscle names: ECR, extensor carpi radialis; ECU, extensor carpi ulnaris; FCR, flexor carpi radialis; FCU, flexor carpi ulnaris; PT, pronator teres; EDC, extensor digitorum communis; FDS, flexor digitorum superficialis (FDSr, FDS radial side); FDP, flexor digitorum profundus (FDPu, ulnar side; FPB, flexor pollicis brevis; FDI, first dorsal interosseous.

**Supplementary Figure2.**
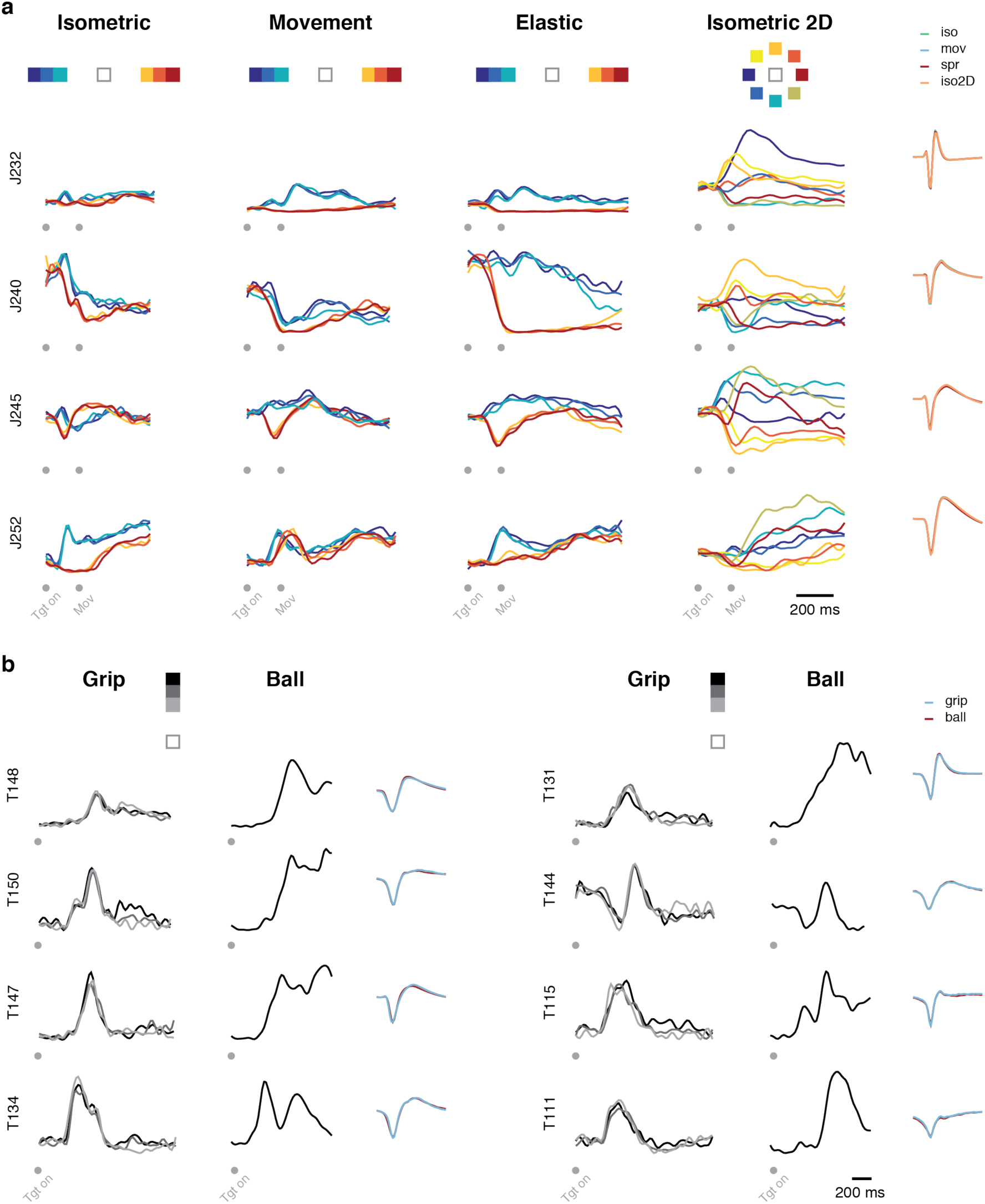
Examples of neural activity patterns illustrate their broad diversity and their complex changes across tasks. **(a)** Peristimulus time histogram (PSTH) of four additional units for one session in which monkey J performed all four wrist tasks (the same tasks as in Fig. 2). PSTHs are colored according to target location (below task name). Right-most columns: mean action potential waveform for each task; each in a different color. **(b)** Peristimulus time histogram (PSTH) of eight units for one session in which monkey T performed the two reach-to-grasp tasks. Data are organized as in (a).

**Supplementary Figure3.**
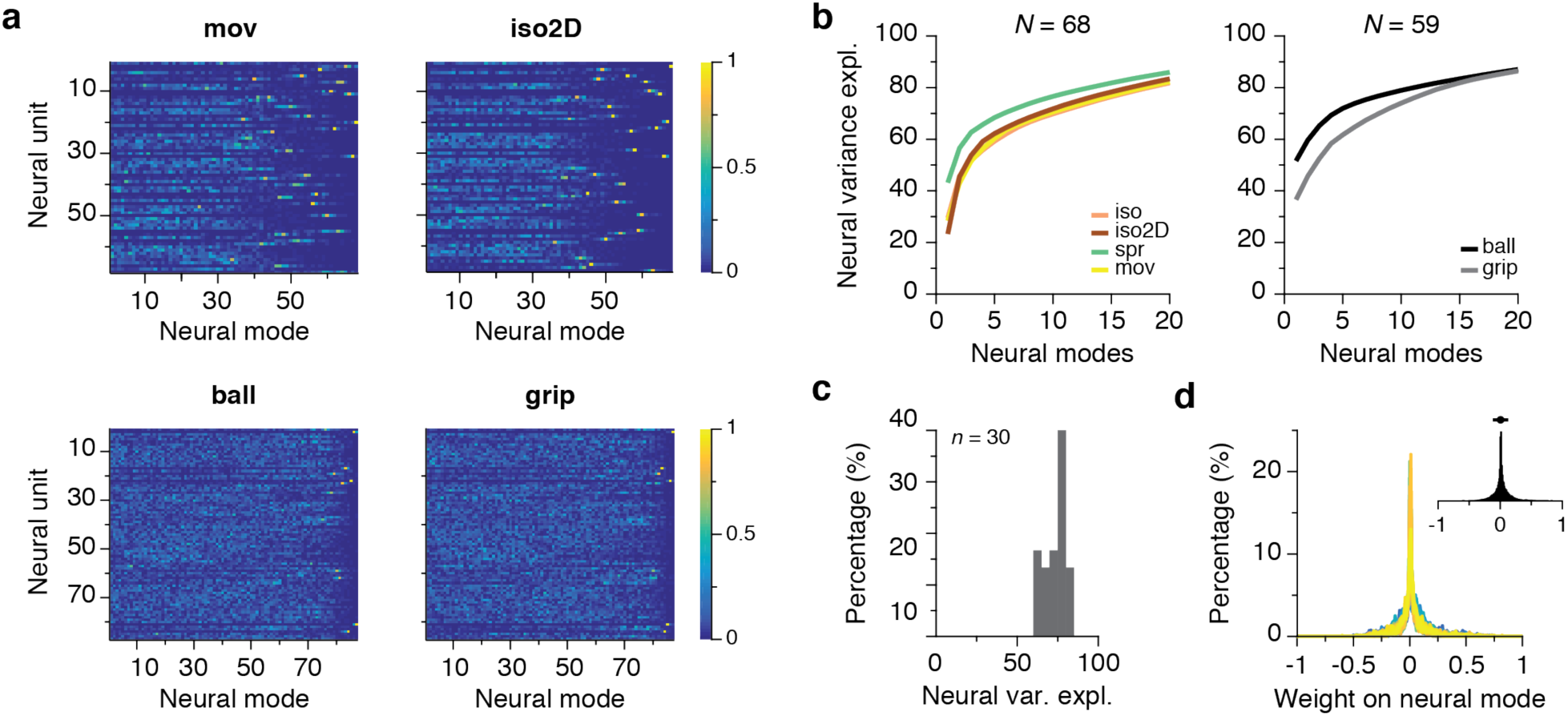
Population dynamics during distal limb tasks are spanned by few neural modes, each involving most units. **(a)** Absolute value of the weights of the neural units onto each PC neural mode for the movement task and two-dimensional isometric task in one session of monkey J (top), and for the ball and grip tasks in one session of monkey C (bottom). Note that most units have weights onto the leading neural modes. **(b)** Neural variance explained as function of the number of neural modes for the four wrist tasks (isometric, movement, spring-loaded movement, and two-dimensional isometric) in one session of monkey J (left), and for the two reach-to-grasp tasks (ball and grip) in one session of monkey T (right). *N*: number of neural units. **(c)** Distribution of neural variance explained by a 12D manifold, pooled over all monkeys, sessions, and tasks. **(d)** Distribution of neural unit weights onto the leading 12 neural modes, across all neural units for each task from each session and monkeys (each shown in a different color). Inset: histogram summarizing all the data (same units as main figure in the panel; error bar: mean ± SD). The units are mostly assigned small weights for all the tasks, and there are no outliers with large weights. The leading neural modes thus do reflect population-wide activity patterns.

**Supplementary Figure4.**
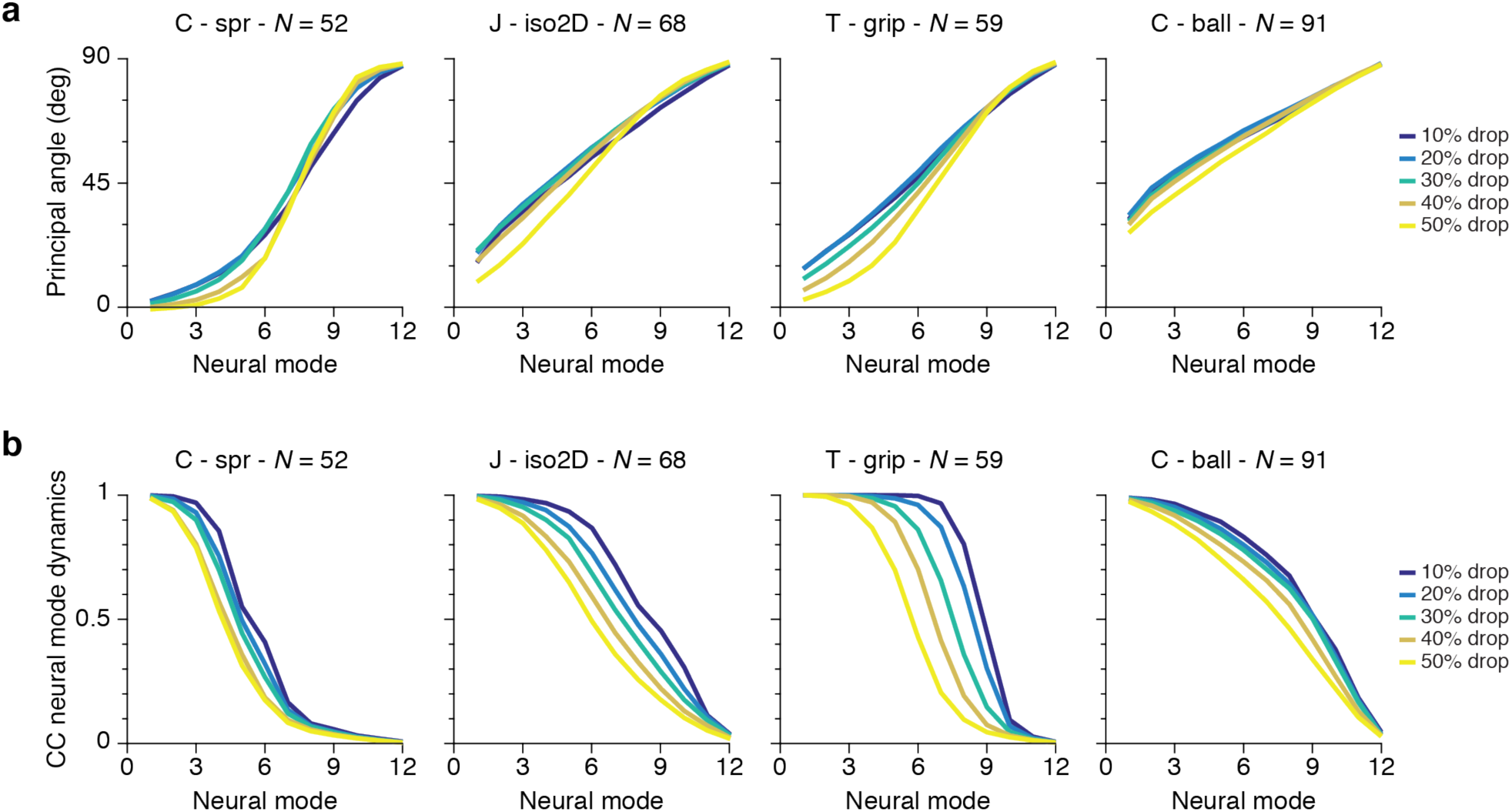
The geometry of the neural manifold and the neural mode dynamics are preserved when dropping a large percentage of units. **(a)** Principal angles between two 12D neural manifolds identified after randomly dropping a given percentage of units (legend). Colored traces: mean principal angle across 100 random drops; title: monkey and task; *N*: number of units. **(b)** Canonical correlation (CC) between the neural mode dynamics computed including all the units and the neural mode dynamics computed after randomly dropping a given percentage of units (legend). Colored traces: mean CC across 100 random drops; title: monkey and task; *N*: number of units.

**Supplementary Figure5.**
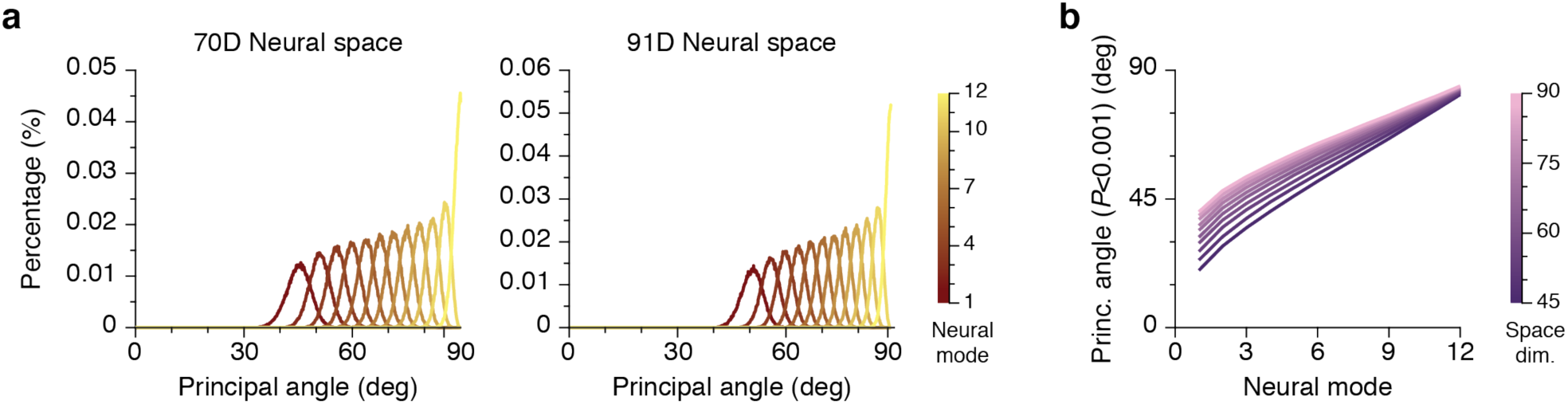
Principal angles between 12D neural manifolds from two different tasks both embedded in a high-dimensional neural space. To interpret the experimentally obtained principal angles, we computed principal angle distributions between pairs of randomly generated manifolds for each neural space dimensionality (the number of units included in each dataset). **(a)** Example distributions of principal angles between 10,000 pairs of 12D randomly generated manifolds in neural spaces with dimensionality *N*=70 and *N*=91. (b) Principal angles that define the *P*< 0.001 significance threshold between pairs of 12D manifolds in neural spaces with dimensionality in the same range as our experimental data. Principal angles below their corresponding significance threshold indicate a large degree of similarity in manifold orientation.

**Supplementary Figure6.**
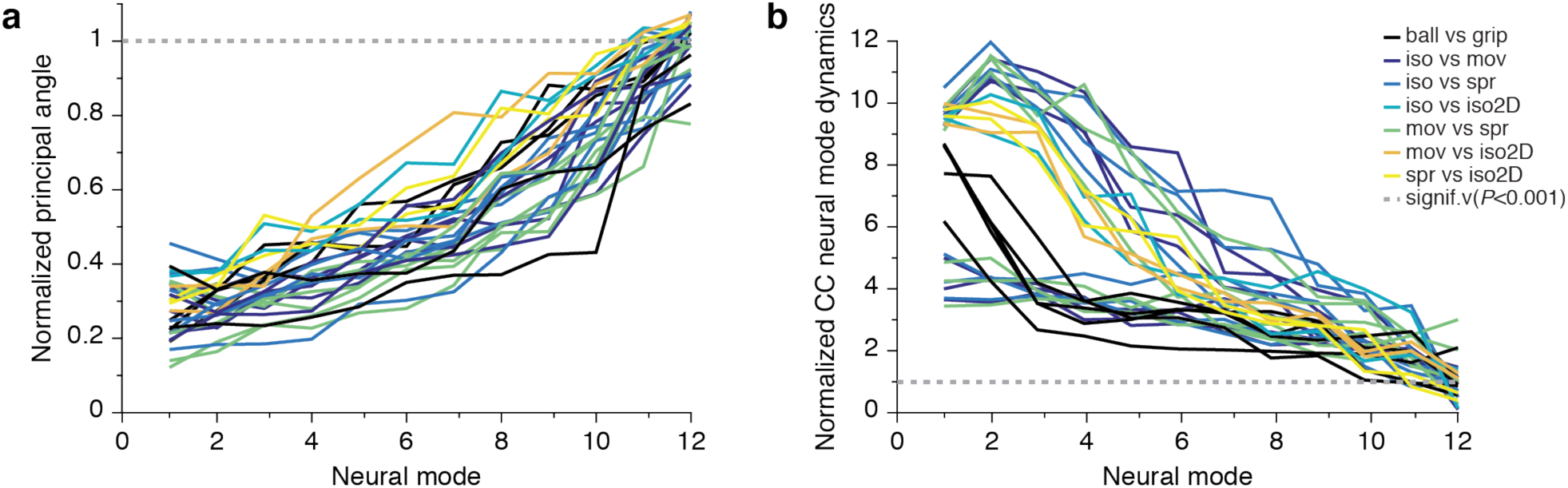
Similarity of the neural modes and their corresponding dynamics across all pairs of tasks. **(a)** Normalized principal angles between the 12D neural manifolds from all pairs of tasks. Data were normalized by dividing the experimentally obtained principal angles by the principal angles that defined the significance threshold (*P*<0.001) for the dimensionality of the corresponding neural space; normalized principal angles <1 are significantly small. The small value of most principal angles indicates that the structure of the neural covariation patters was well preserved across wrist and reach-to-grasp tasks. **(b)** Normalized canonical correlation (CC) between the neural mode dynamics from all pairs of tasks. Similar to (a), data were normalized by dividing the experimentally observed CCs by the CCs that defined the significance threshold (*P*< 0.001). Therefore, CCs >1 indicate that the neural mode dynamics were significantly similar. Many of the leading CCs were well above the significance threshold, suggesting that the dynamics of several neural modes were well preserved across motor tasks.

**Supplementary Figure7.**
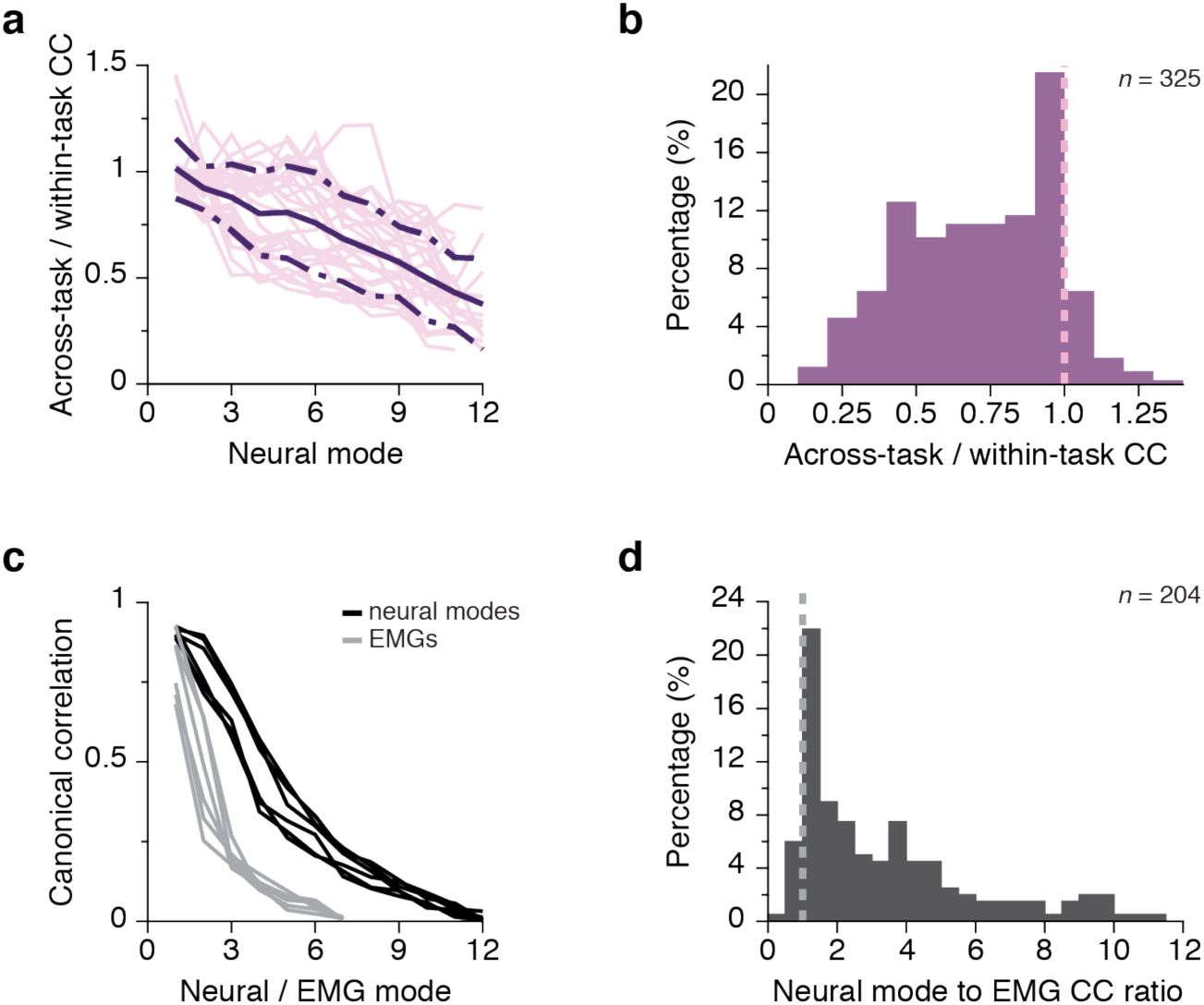
Similarity of neural mode dynamics across tasks and comparison between across-task neural mode dynamics and EMGs. **(a)** Ratio of the across-task CC between neural mode dynamics to the within-task CC between neural mode dynamics. Data for all monkeys, sessions, and task comparisons (pink traces: individual comparisons; purple traces: mean ± s.d.). Ratios < 1 confirm that within-task correlations provide an upper bound to across-task correlations. The leading ratios are quite large (the average ratio for the leading 6 dimensions was always ≥ 0.75), indicating a remarkably high across-talk correlations between leading neural modes. Only the ratios for projections with significant across-task correlation were included. **(b)** Summary of the data in (a), pooled across all manifold dimensions. **(c)** Canonical correlation between the neural mode dynamics from six pairs of wrist tasks compared to the canonical correlations between the corresponding muscle activation patterns (EMGs). Data are the same as in Fig. 4c (right). For this representative example, the neural mode dynamics were more preserved than the muscle activity patterns, overall for dimensions 2–7. **(d)** Ratio of the across-task CC between neural mode dynamics to the across-task CC between EMGs (as shown in (c)), pooled over all tasks, sessions, and monkeys. Most values are >1 (dashed vertical line), indicating that the structure of the neural mode dynamics was in general better preserved across tasks than the structure of the EMGs.

**Supplementary Figure8.**
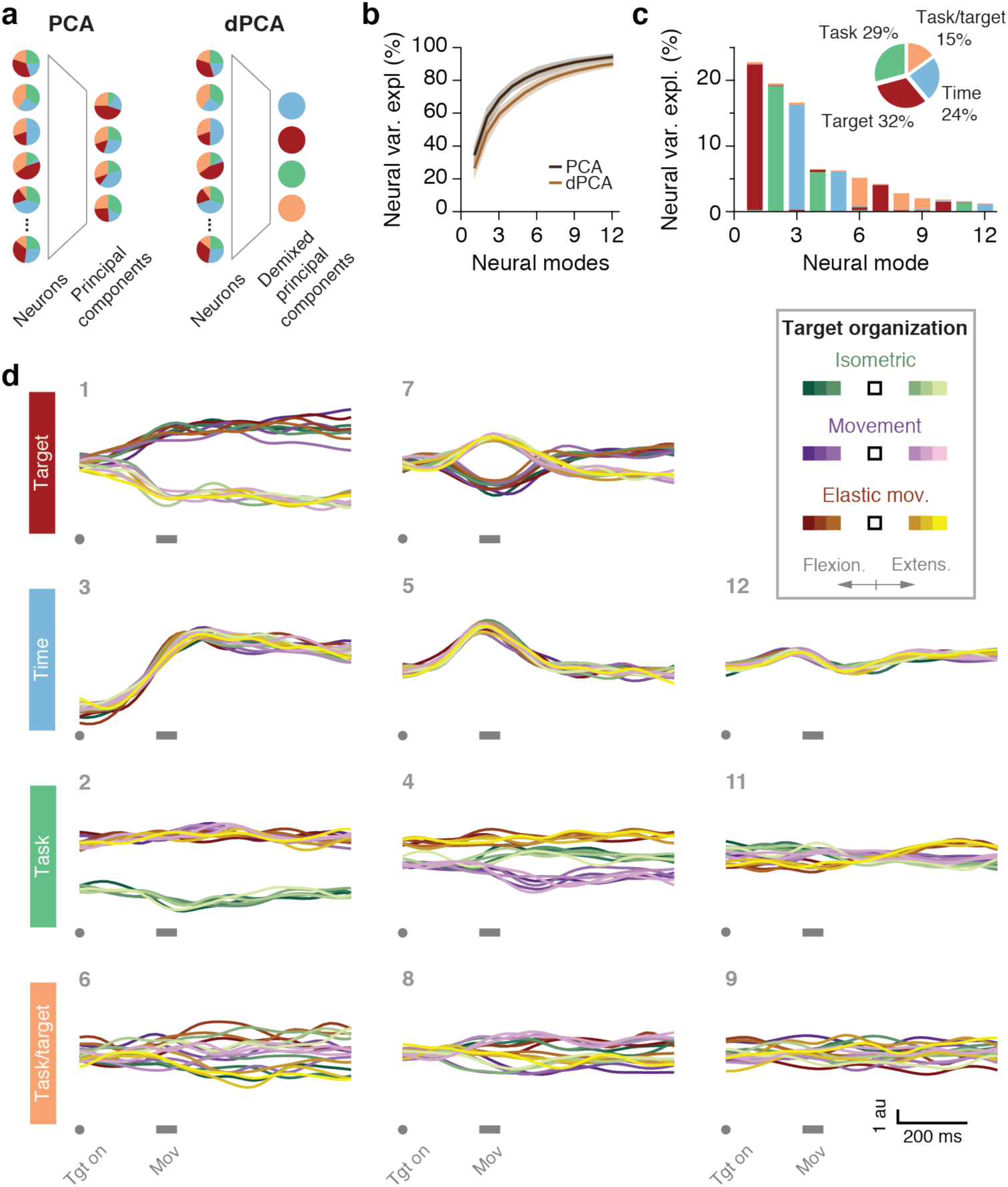
Additional information on task-specific and task-independent neural mode dynamics identified by dPCA. **(a)** Dimensionality reduction with PCA and dPCA. Unlike PCA, dPCA identifies neural modes that are linear readouts of activity associated with relevant behavioral parameters. **(b)** Neural variance explained by 12D manifolds spanning all the wrist tasks from each session, identified with either PCA or dPCA (legend). Plot shows mean ± s.d. (trace and colored strip). **(c)** Neural variance explained by each dPC neural mode, and its relation to behavioral parameters for one example session from monkey C. Inset: amount of neural variance associated with each behavioral parameter across all twelve modes. **(d)** Activation dynamics of eleven neural modes, grouped in four sets based on the behavioral parameter they are most strongly associated with. The number on the top left of each panel indicates the ranking of that neural mode in terms of neural variance explained, as in (c). Each row corresponds to one behavioral parameter (vertical labels on the left). Each panel has 18 traces, corresponding to each of the 18 task-target combinations (bottom inset: target locations for each task and color code for each task-target combination. Extension targets are shown in dark colors, flexion targets in light colors; each task is represented using a different color). Panel (a) adapted from Ref. 36.

**Supplementary Figure9.**
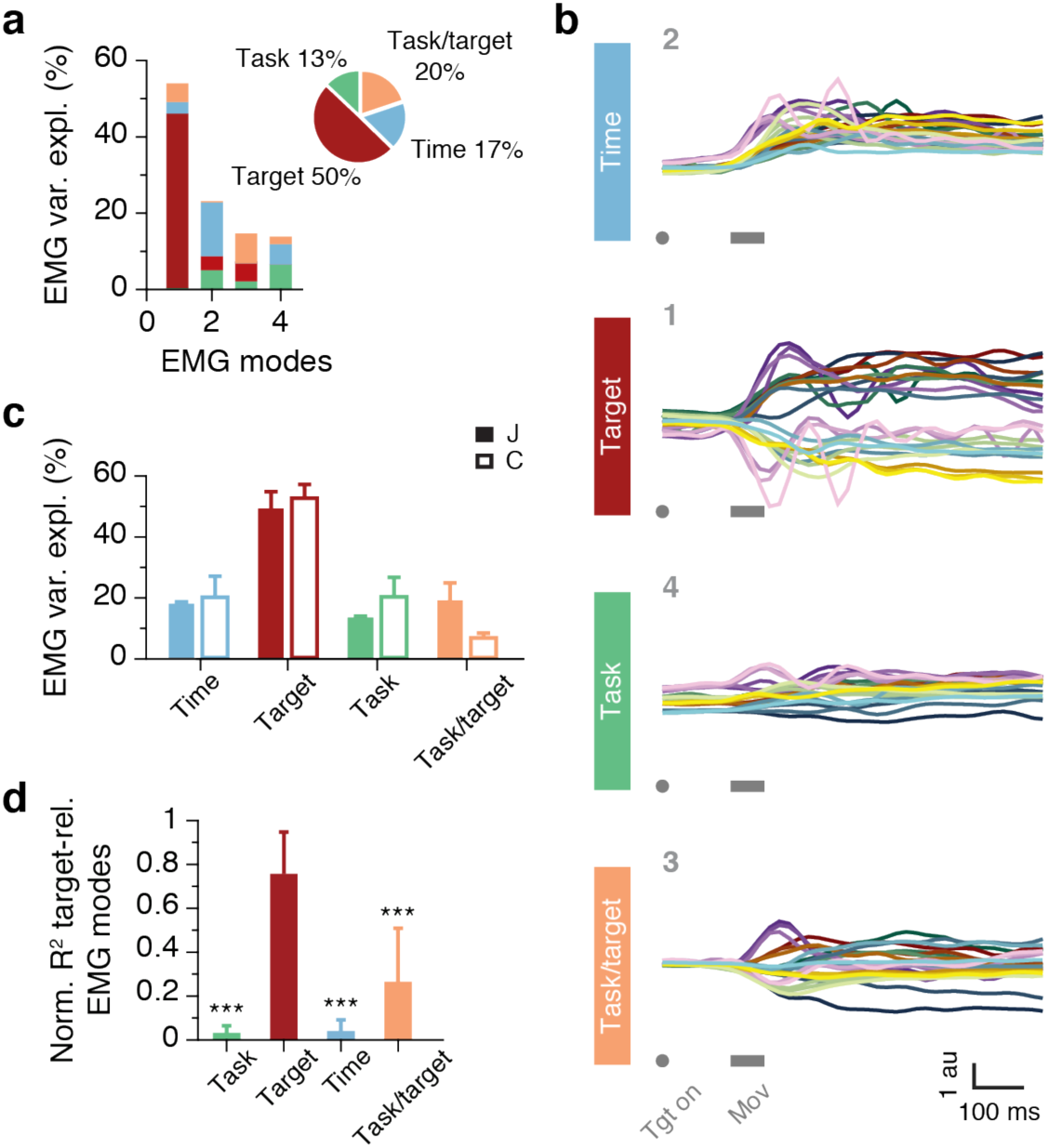
Decomposition of the muscle activity (EMG) associated with the wrist tasks into EMG modes. **(a)** EMG variance explained by each EMG mode, and its relation to behavioral parameters for the example session from monkey J shown in Fig. 5. Inset: amount of neural variance associated with each behavioral parameter across all four modes. **(b)** Activation dynamics of the four EMG modes, grouped in four sets based on the behavioral parameter they are most strongly associated with. The number on the top left of each panel indicates the ranking of that neural mode in terms of EMG variance explained, as in (a). Each row corresponds to one behavioral parameter (vertical labels on the left). Each panel has 24 traces, corresponding to each of the 24 task/target combinations; color code shown in Fig. 5. **(c)** Total amount of EMG variance explained for each behavioral parameter, averaged for all the sessions from monkeys J and C. Bars: mean + s.d. **(d)** Normalized R^2^ of the predictions of the target-related EMG modes obtained from four types of decoders; each type used as inputs the activations of the two leading neural modes most strongly related to each of the four dPCA behavioral parameters. Performance was averaged over all muscles, tasks, and monkeys. Bars: mean + s.d; the *** denotes *P*∼ 0 (paired *t*-test).

